# A positively Tuned Voltage Indicator Reveals Electrical Correlates of Calcium Activity in the Brain

**DOI:** 10.1101/2021.10.21.465345

**Authors:** S. Wenceslao Evans, Dongqing Shi, Mariya Chavarha, Mark H. Plitt, Jiannis Taxidis, Blake Madruga, Siri C. van Keulen, Michelle M. Pang, Sharon Su, Fuu-Jiun Hwang, Guofeng Zhang, Austin Reese, Lagnajeet Pradhan, Jiang Lan Fan, Sungmoo Lee, Yu Liu, Carl-Mikael Suomivuori, Dongyun Jiang, Adrian Negrean, Sui Wang, Na Ji, Thomas R. Clandinin, Ron O. Dror, Guoqiang Bi, Christopher D. Makinson, Peyman Golshani, Lisa M. Giocomo, Attila Losonczy, Jun B. Ding, Michael Z. Lin

**Affiliations:** Department of Neurobiology, Stanford University Medical Center, Stanford, CA, USA; School of Life Sciences, University of Science and Technology of China, Hefei, China; Department of Bioengineering, Stanford University, Stanford, CA, USA; Department of Neuroscience, Columbia University, New York, NY, USA; Mortimer B. Zuckerman Mind Brain Behavior Institute, New York, NY, USA; Department of Neurology, UCLA David Geffen School of Medicine, Los Angeles, CA, USA; Department of Neurosurgery, Department of Neurology and Neurological Sciences, Stanford University Medical Center, Stanford, CA, USA; Department of Computer Science, Stanford University, Stanford, CA, USA; Departments of Structural Biology and of Molecular and Cellular Physiology, Stanford University School of Medicine, Stanford, CA, USA; Institute for Computational and Mathematical Engineering, Stanford University, Stanford, CA, USA; Department of Ophthalmology, Stanford University Medical Center, Stanford, CA, USA; UC Berkeley/UCSF Joint Program in Bioengineering, University of California Berkeley, Berkeley, CA; Department of Molecular and Cell Biology, University of California Berkeley, Berkeley, CA; Department of Physics, University of California Berkeley, Berkeley, CA; Helen Wills Neuroscience Institute, University of California Berkeley, Berkeley, CA; Molecular Biophysics and Integrated Bioimaging Division, Lawrence Berkeley National Laboratory, Berkeley, CA, USA; Department of Neurosurgery, Stanford University Medical Center, Stanford, CA, USA; Kavli Institute for Brain Science, New York, NY, USA

## Abstract

Neuronal spiking activity is routinely recorded using genetically encoded calcium indicators (GECIs), but calcium imaging is limited in temporal resolution and does not report subthreshold voltage changes. Genetically encoded voltage indicators (GEVIs) offer better temporal resolution and subthreshold sensitivity, but spike detection with fast GEVIs has required specialized imaging equipment. Here, we report the ASAP4 subfamily of genetically encoded voltage indicators (GEVIs) that brighten in response to membrane depolarization, inverting the fluorescence-voltage relationship of previous ASAP GEVIs. Two variants, ASAP4b and ASAP4e, feature 128% and 178% fluorescence increases over 100-mV of depolarization, respectively, facilitating spike detection in single trials *in vivo* with standard 1 and 2-photon imaging systems. Simultaneous voltage and calcium imaging confirms improved temporal resolution and spike discernment by ASAP4 GEVIs. Thus, positively tuned ASAP4 voltage indicators enable recording of neuronal spiking activity using similar equipment as calcium imaging, while providing higher temporal resolution.

**One Sentence Summary:** Upward ASAPs increase detection capability of GEVIs *in vivo*.

## INTRODUCTION

Recording the real-time activity of genetically specified neurons during animal behavior will be crucial for understanding how the nervous system relays and processes information (*3, 48*). Currently, most genetically targeted neuronal activity recording is performed with genetically encoded calcium indicators (GECIs), which can produce large persistent signals that are easily discerned with either 1-photon or 2-photon imaging. Detection of calcium activity is aided by high cytosolic expression levels of GECIs, low fluorescence at basal calcium levels, the long duration of calcium transients, and the ability of calcium to increase its cytosolic concentration non-linearly during bursts of action potentials (APs), dramatically increasing the fluorescence of the GECI during those events. However, GECI signals lack precise temporal resolution of the timing of spikes, as somatic calcium continues rising after the peak of the action potential, due to delayed inactivation of voltage-gated calcium channels and continued diffusion of calcium from membrane influx points into the cytosol (*16, 45*). This temporal smoothing effect, together with >200-ms half-extrusion times for calcium, prevents GECIs from accurately discerning closely spaced APs (*48*). Lastly, calcium levels do not respond appreciably to subthreshold and hyperpolarizing changes in transmembrane voltage and thus do not reveal subthreshold dynamics such as theta rhythms and hyperpolarizing events, as these events do not generally open voltage activated calcium channels.

Imaging GEVIs can provide more accurate spike timing information while also revealing subthreshold dynamics, but suffers from signal detection challenges. The membrane localization and temporal transience of voltage signals present obstacles unique to voltage imaging. Unlike GECIs which can detect calcium changes from the cytosol, GEVIs must exist at the membrane to report voltage changes, resulting in fewer molecules and thus fewer photons per imaging volume (*6*). Electrical events are also much faster than calcium events, and GEVIs that track electrical kinetics produce similarly short-lived signals. This limits the number of integrated photons per transient, and requires fast sampling rates to detect the response. For example, the GEVI ASAP3 produces a similar relative fluorescence change (Δ*F*/*F*_0_) to a single AP as the widely used GECI GCaMP6 (∼20%), and actually does so with ∼20-fold higher per-molecule brightness (*39*). However, its expression level per cell body is estimated to be ∼20-fold less, and its signal is 20-fold shorter in duration (*39*). To combat these formidable challenges, GEVI engineering over the last several decades has focused on increasing the per-molecule change in fluorescence during electrical activity while optimizing kinetics. In particular, brighter fluorophores and fast onset kinetics allow the largest modulation in absolute photons per molecule, while slightly slower offset kinetics improve detectability by allowing the photon flux change to persist for longer.

Currently, the GEVIs with the largest relative response per AP are the opsin-based GEVI Archon1 and the voltage sensing domain (VSD)-based GEVI ASAP3. Archon1 consists of a non-conducting opsin domain and demonstrates voltage-dependent fluorescence upon red light illumination, due to shifting of absorbance by voltage-dependent protonation of the retinal Schiff base. Unfortunately, their inherent fluorescence is dim, requiring high intensity illumination to detect (*29*). Opsins can be fused to a brighter fluorophore, allowing the fluorophore brightness to be modulated by FRET in a voltage-dependent manner, but the degree of modulation is limited (*1, 2, 14, 23*). Notably, both types of opsin-based GEVIs show poor responsivity under 2-P illumination, likely because light absorption is required to enter the voltage-sensitive fluorescent state, but this state decays too quickly for the pulsed 2-P lasers to interrogate (*6, 8, 24*).

In contrast to opsin-based GEVIs, ASAP3 is composed of a circularly permuted GFP (cpGFP) inserted in an extracellular loop of a VSD (*35, 39*). Voltage-dependent movements of the VSD are then transduced to alter cpGFP fluorescence, analogous to cpGFP-based indicators such as GCaMP (**Fig. S1A)**. ASAP3 features a large response range of 51% from -70mV to +30mV, is reasonably bright, sufficiently fast enough to track trains of action potentials at up to 100hz on single trials, and can be used in awake and behaving flies and mice (*8, 39, 45*). (*8, 39, 45*)While it may be surpassed in individual parameters of kinetics, relative responsiveness, or per-molecule peak brightness by different indicators, ASAP3 performs well across multiple parameters and thus exhibits the highest power-normalized signal-to-noise ratio for both spiking and subthreshold activity (*26*). In addition, ASAP-family GEVIs operate equally well under 1-P and 2-P illumination (*8, 39*). We set out to improve ASAP-family GEVIs in SNR and versatility. An important goal in GEVI engineering is to create indicators with low baseline fluorescence that brighten with positive voltage changes. Regardless of directionality, the largest possible change in signal for a fluorescent indicator is determined by the molar brightness of the fluorophore. However, in the shot noise-limited regime, noise is related to the square root of baseline fluorescence, and thus an indicator with low baseline fluorescence would have the benefit of lower baseline noise, while maintaining higher SNR during depolarizing events as the ratio of 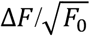 increases. A well engineered positively tuned GEVI would have the ability to surpass all negatively tuned GEVIs in SNR (*2*). A positively tuned GEVI also allows experimenters to minimize long-term photodamage when imaging the same cells in vivo over many days by setting laser power to the minimum necessary to observe baseline fluorescence. Among previously described fully genetically encoded voltage indicators with high fluorescence, only ElectricPk, FlicR1, Marina and upward Ace2N-mNeon are positively tuned. Like ASAPs, ElectricPk and FlicR1 are based on fusions of a VSD with a circularly permuted fluorescent protein. Marina also uses a VSD, but attaches a mutated version of the pHluorin fluorescent protein to the C-terminus at the end of the S4 helix. Upward Ace2N-mNeon is based on electrochromic FRET between the mNeonGreen fluorescent protein and a voltage-sensing opsin (*2*). ElectricPk, FlicR1 and Marina responses to 100-mV steps were reported as 1.2%, 2.5% and 29.2% (*2, 4, 26*), but Ace2N-mNeon responses to 100-mV steps has not been reported, although it appeared to report APs with larger fluorescence changes than ElectricPk and FlicR1 (*2*).

Here, using PCR transfection, electroporation, and computational protein modeling (*39, 41*), we develop two new GEVIs, ASAP4b and ASAP4e, that brighten in response to membrane depolarization. One variant, ASAP4e, features a steady-state voltage change of 178% over the physiological range and 468% over all voltages, while another variant, ASAP4b, combines faster activation with higher basal and peak brightness. We demonstrate that ASAP4e and ASAP4b report APs and sub-threshold oscillations using standard one-photon (1-P) or two-photon (2-P) equipment, in awake behaving animals. By simultaneously imaging red GECIs, we verify that ASAP4 GEVIs provide higher temporal resolution with better SNR for single APs.

## RESULTS

### Engineering of positively tuned GEVIs

Our goal was to create fluorescent GEVIs that respond positively, i.e. increase in brightness with positive (depolarizing) changes in the transmembrane potential. Positively tuned GEVIs have a major theoretical advantage in potentially achieving higher SNR than negatively tuned GEVIs. In the shot-noise limited regime, where photons are Poisson distributed, SNR is proportional to 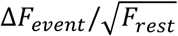, where Δ*F*_*event*_ is the change in photons emitted from the imaged population of molecules due to the event of interest per time interval, and *F*_*rest*_ is photons emitted per time interval at rest. With a positively tuned GEVI, it is theoretically possible to achieve an *F*_*rest*_ near zero while having Δ*F*_*event*_ approaching the entire maximum brightness of the fluorophore population. However, previous ASAP-family GEVIs brighten upon negative (hyperpolarizing) voltage changes.

As with most other GFP-based indicators, ASAP-family GEVIs are imaged by excitation with ∼490-nm light in 1-P mode or ∼940-nm light with 2-P illumination. This wavelength selectively excites the deprotonated anionic form of the GFP chromophore with peak absorbance ∼490 nm, avoiding the protonated or neutral GFP chromophore with peak ∼390-nm absorbance (*38, 40*). In wild-type GFP, hydrogen bond donation by His-148 (His-151 in previous ASAP variants) stabilizes the deprotonated state. We hypothesized that depolarization-induced movements in the S4 transmembrane helix cause ASAP dimming by disrupting hydrogen bond donation by His-148. This made His-151 and the neighboring Ser-150 (**Fig. S1B)** excellent candidates for inverting the relationship between S4 movement and chromophore protonation.

Starting with ASAP2f L146G S147T R414Q, an intermediate in ASAP3 evolution (*39*), we constructed and tested all 400 possible combinations of amino acids at positions 150 and 151 by 384-well PCR transfection followed by automated electroporation and imaging (*39*). We identified several mutants with upward responsivity, one of which, S150D H151G (ASAP4.0), demonstrated a 2-fold increase in fluorescence upon voltage steps from –70 mV to +30 mV (**Fig. S1C, S2**). However, ASAP4.0’s fluorescence-vs-voltage (*F-V*) curve was right-shifted, such that the physiological voltage range of –70 to +30 mV mapped to the shallow lower portion of the *F-V* curve (**Fig. S1D, S1E**). As a result, the fluorescence change from –70 to +30 mV (Δ*F*_100_) was only 29% of the maximum obtainable fluorescence across all voltages (*F*_*max*_), and the fluorescence at +30 mV (*F*_+30_) was less than half of *F*_*max*_ (**Fig. S1E**). In addition, ASAP4.0 responses demonstrated partial reversion during depolarization steps (**Fig. S1D**), suggesting a relaxation process that reduces the proportion of activated molecules and/or shifts a proportion to a dimmer conformation.

Over 4 rounds of structure-guided mutagenesis in the GFP and linker regions, we were able to left-shift and sharpen the curve (details in **Supplementary Note 1**), producing large responses in the physiological voltage range (**Table S1**). To reduce the rapid relaxation and obtain higher *F*_+30_*/F*_*max*_, we mutated a chromophore-interacting site in ASAP4.0 with the goal of stabilizing the bright state of the chromophore. We obtained a mutant, ASAP4.1, with a brighter activated state (**Fig. S3A)**, which also exhibited less relaxation (**Fig. S1D**). However, ASAP4.1 also showed higher fluorescence at baseline (*F*_–70_*/F*_*max*_), reducing the relative response size to steady-state 100-mV depolarizations (Δ*F*_100_*/F*_*–70*_, **Fig. S1E**). We then performed double-site saturation mutagenesis at positions F148 and N149 in the linker between the third transmembrane helix of the VSD (S3) and cpGFP **(Fig. S3B)**, obtaining ASAP4.2 with F148P N149V, with a modestly improved Δ*F*_100_*/F*_*–70*_ (**Fig. S1D**), due to lower *F*_*–*70_ (**Fig. S1E**). Finally, by developing an atomic model of ASAP4 (*34*), we identified Phe-413 as a residue at the end of the fourth transmembrane helix (S4) that may modulate the energetic barrier to voltage-dependent movements of S4 (**Fig. 1A**) (*15*). Mutation of Phe-413 produced several mutants with larger Δ*F*_100_*/F*_-70_ (**Fig. S4A)**.

**Figure 1.**
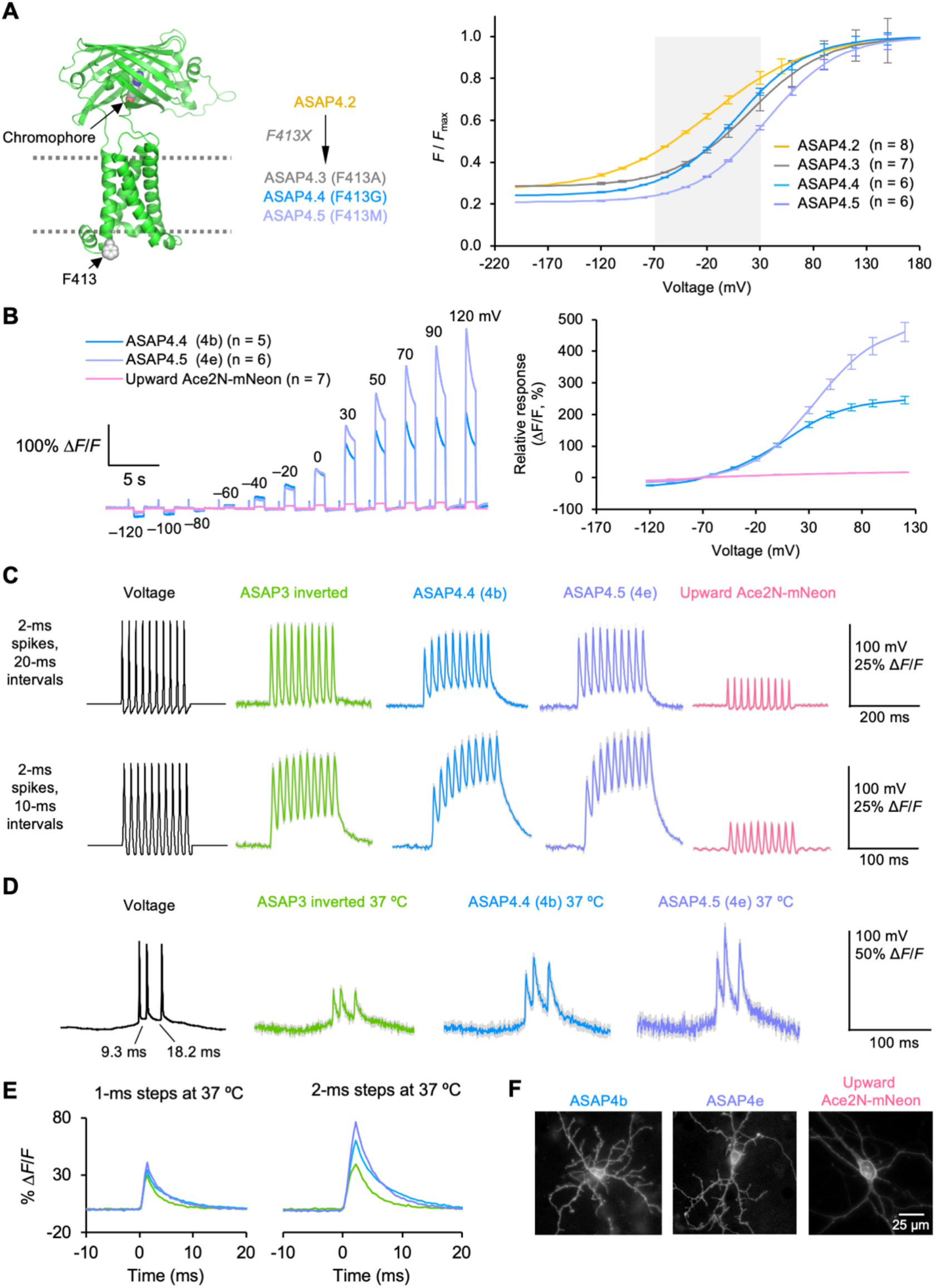
The ASAP4-family of positively tuned GEVIs. **(A)** ASAP4.2 model with position 413 shown in gray and mutations in final ASAP4 variants. Right, F-V curves for ASAP4.2 to ASAP4.5 fit to a sigmoid curve, normalized to maximum fluorescence. Error bars, SEM. **(B)** Left, steady-state responses to voltage commands in HEK293A cells of ASAP4.4 (4b), ASAP4.5 (4e), and upward Ace2N-mNeon, normalized to –70mV. Right, resulting F-V curves. Error bars, SEM of n = 5 cells for ASAP4b-Kv, 6 cells for ASAP4e-Kv, and 7 cells for upward Ace2mNeon for both figures. **(C)** Response to commanded spike trains in HEK293A cells at room temp. V_m_ was –70 mV at the start of the displayed segment and peak AP amplitude was 100 mV. Responses were collected from 6 (ASAP4.2), 7 (ASAP4b), 5(ASAP4e), 6 (ASAP3) and 7 (Upward Ace2mNeon) HEK293a cells, each supplying a single unfiltered and unsmoothed 1000-Hz trace. Colored lines represent averages, gray shading is SEM. **(D)** Responses of indicators in HEK293 cells to an AP burst waveform with spike widths of 1.0 ms at FWHM, previously recorded from the mouse hippocampus at 37 ºC. V_m_ was –52 mV at the start of the displayed segment. Each colored line is the mean of 5 HEK293 cells, each supplying a single unfiltered and unsmoothed trace. Gray shading, SEM. **(E)** Left, average responses to a 1-ms square voltage step from –70 to +30mV in HEK293A cells at 37º C. Right, similar, but for a 2-ms pulse. The number of pulses was 50 (ASAP3), 50 (ASAP4b) and 60 (ASAP4e). **(F)** Example images of ASAP4b, ASAP4e, and upward Ace2N-mNeon expressed in cultured rat hippocampal neurons. ASAP4b and ASAP4c brightness was scaled 10-fold relative to upward Ace2N-mNeon.

Among these mutants, ASAP4.4 and ASAP4.5 had the steepest F-V curves (**Fig. S4B, 1A)**, resulting in Δ*F*_100_*/F*_*–*70_ of 128% and 178% respectively (**Fig. 1B**). Ace2N-mNeon was previously the most responsive positive indicator based on responses to APs, but no published F-V curve exists for it (*2*). We measured its responses for comparison, obtaining Δ*F*_100_*/F*_–70_ of 12% (**Fig. 1B**). ASAP4.4 and ASAP4.5 report physiological voltage changes with fluorescence changes that are larger than previously described positively tuned GEVIs (*30*). Given their optical characteristics, we named ASAP4.4 as ASAP4b (brighter baseline) and ASAP4.5 as ASAP4e (enhanced response).

Since all ASAP4 variants after ASAP4.2 share an identical GFP fluorophore, their maximum obtainable fluorescence per molecule should be identical. Fluorescence at any given voltage as a fraction of maximal fluorescence across all voltages in the same patch-clamp experiment should reflect per-molecule brightness at that voltage. Between ASAP4b and ASAP4e, ASAP4b had the higher fractional brightness at both –70 mV and +30 mV, while ASAP4e had the larger change relative to baseline due to its lower baseline brightness (**Fig. 1A**). Both reached >50% of peak fluorescence at +30 mV. While previous studies obtained steady-state F-V curves for GEVIs at room temperature only, we also characterized F-V relationships for ASAP3, ASAP4b, and ASAP4e at 37 ºC. Interestingly, the F-V curves of ASAP4b and ASAP4e steepened and became more similar, due to lower *F*_*–*70_*/F*_*max*_ for ASAP4b and higher *F*_+30_*/F*_*max*_ for ASAP4e, while the F-V curve for ASAP3 underwent a left-shift (**Fig. S4C**).

To obtain preliminary confirmation that ASAP4-family GEVIs can function well *in vivo*, we performed 2-P imaging of ASAP4b in the axon termini of L2 neurons in the fruit fly visual system during visual stimulation with a contrast stimulus. These neurons respond bidirectionally to luminescence changes, with membrane hyperpolarization upon increased luminescence, and depolarization upon decreased luminescence. We observed ASAP4b fluorescence decreased in response to a transient bright stimulus, hyperpolarizing L2, and increased in response to a dark stimulus, depolarizing L2, opposite in direction to the negatively tuned indicator ASAP2f (**Fig. S5**), confirming the inverted voltage response of ASAP4b. Response amplitudes were approximately 2-fold higher for ASAP4b than ASAP2f to stimuli in either direction. Activation kinetics of the two indicators were similar. These results indicate that ASAP4b can be used to image subcellular electrical activity in vivo, and improves upon the response properties of ASAP2f in flies.

### Characterization of ASAP4 kinetics, photostability, and brightness

To understand how well ASAP4b and ASAP4e can report voltage dynamics of different speeds, we measured the kinetics of fluorescence responses to voltage steps at room temperature and 37 ºC. At room temperature, activation kinetics were well modelled as a sum of two exponentials. In response to depolarization, ASAP4e was faster than ASAP4b, with fast activation kinetics of 2.6 ms vs 3.9 ms, accounting for 14% and 19% of the response amplitude, respectively (**Table S2**). At 37 ºC, ASAP4b and ASAP4e activation kinetics accelerated, with the fast component reaching similar values of 1.5 ms each (**Table S2)**.

We next tested responses to AP waveforms. At room temperature, responses to APs followed the rank order ASAP4b < ASAP3 < ASAP4e, with 19%, 22%, and 23% fluorescence changes to APs of 2-ms full-width at half-maximum (FWHM) (**Fig. 1C)**. In response to 4-ms APs, which can be more easily compared to previously reported results (*29*), fluorescence changes were 26%, 29%, and 34%, respectively (**Fig. S6A**). In contrast, upward Ace2N-mNeon (*2*) showed 8% or 10% responses to 2-ms or 4-ms APs, respectively. All GEVIs membrane localized well in cultured rat hippocampal neurons (**Fig 1F**), and were able to discern single spikes within 50- and 100-Hz bursts of APs. The presence of a slowly activating component and slower deactivating kinetics compared to ASAP3 are apparent in ASAP4b and ASAP4e (**Fig. 1C)**.

We next tested responses to naturalistic AP waveforms at 37 ºC in HEK293 cells by voltage clamp. Response amplitudes to imposed voltage waveforms derived from mouse hippocampal pyramidal neuron recordings (baseline V_m_ of –60 mV, AP FWHM of 1 ms) followed the rank order ASAP3 < ASAP4b < ASAP4e (**Fig. S6B-S5E**). Responses to single APs were 24%, 30%, and 41%, respectively (**Fig. S6B**). Individual spikes within a triplet burst with interspike intervals of 9.3 and 18.2 ms were well discerned (**Fig. 1D**). The higher fluorescence reached by ASAP4b and ASAP4e, but not ASAP3, upon the second spike in the burst is expected from the slowly activating component in ASAP4b and ASAP4e (**Table S2**). This becomes more apparent as the stimulus duration increases, giving ASAP4 more time to take advantage of its massive dynamic range (**Fig. 1E**).

We next compared photostability under 1-P or 2-P illumination for ASAP4b, ASAP4e, and ASAP3 in cultured cells. Under 453 nm 1-P illumination at 50 mW/mm^2^, ASAP3 exhibited mono-exponential photobleaching while ASAP4e and ASAP4b increased in brightness for several minutes before finally decaying (**Fig. S7A, Table S3**). ASAP4e and ASAP3 were more photostable than ASAP4b during 940-nm 2-P illumination (**Fig. S7B, Table S3**). Most of the photobleaching represented reversibly photoswitchable events, as most of the fluorescence reduction was reversed by a few minutes of incubation in the dark.

As a metric of relative baseline fluorescence per molecule, *F*_-70_/*F*_*max*_ measured in HEK293 cells followed the order ASAP4.2 > ASAP4.3 ∼ ASAP4b > ASAP4e (**Table S1**). However, actual per-molecule brightness at –70 mV may not follow the same trend if specific variants mature more poorly. In addition, the T206H mutation introduced in ASAP4.1 interacts directly with the GFP chromophore and slightly red-shifts it (*20*), and could thus alter the maturation or molecular brightness of the subsequent ASAP4 variants compared to ASAP3. To empirically determine relative per-molecule brightness of somatically targeted ASAP variants in neurons, we fused ASAP1, ASAP3, and each ASAP4 variant to a Kv2.1 segment that we had established to efficiently target ASAP GEVIs to somata *in vivo* (*10, 39*), followed by the cyan-excitable red fluorescent protein mCyRFP2 which we recently developed to provide a non-aggregating voltage-independent label for ASAP GEVIs (*5*). Quantification of the green/red fluorescent ratio upon cyan light excitation found that resting brightness followed the trend ASAP3-Kv > ASAP1-Kv > ASAP4.3-Kv ∼ ASAP4b-Kv > ASAP4e-Kv = ASAP4.2-Kv (**Fig. S7C**). Other than ASAP4.2-Kv, whose low per-molecule brightness despite a high *F*_-70_/*F*_*max*_ ratio suggests poor chromophore maturation, each GEVI produced relative per-molecule brightness values in the rank order expected from their *F*_-70_/*F*_*max*_ ratios. Note that, as SNR scales with 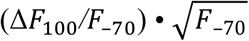, the increased molecular brightness of ASAP4b compensates partially for its relative response being lower than ASAP4e (**Tables S1**).

Taken together, our measurements indicate that ASAP4b and ASAP4e have the largest fluorescence responses of any GEVIs described so far, both relative to baseline and in absolute brightness change per molecule, and for both slow steps and fast spikes (**Tables S1)**.

### ASAP4 expression and performance in mammalian brain tissue

To test the new GEVIs in genetically defined neuronal populations in the mouse, we prepared an adeno-associated virus (AAV) with cre-dependent expression elements to express ASAP4b-Kv, ASAP4e-Kv, or ASAP3-Kv. We first examined expression in medium spiny neurons of the dorsal striatal medium 28-36 days after viral injection. All three GEVIs could easily be observed with simple 1-P illumination, and were expressed predominantly at the plasma membrane (**Fig. S8A**). In terms of whole-cell fluorescence, ASAP3-Kv appeared slightly brighter than ASAP4b-Kv and ASAP4e-Kv, although the differences were not significant (**Fig. S8B**). Given the approximately two-fold higher molecular brightness of ASAP3-Kv at rest (**Fig. S7C**), this result suggested similar or better expression of ASAP4b-Kv and ASAP4e-Kv in these neurons *in vivo* compared to ASAP3-Kv. In hippocampal somatostatin-positive (SST+) interneurons, ASAP4b-Kv and ASAP4e-Kv were approximately half as bright as ASAP3 at rest (**Fig. S8C**), suggesting similar expression of all three GEVIs. We also compared the brightness of ASAP4b-Kv to ASAP3-Kv under 2-P illumination in hippocampal pyramidal neurons. Per-cell baseline fluorescence trended higher for ASAP3-Kv than ASAP4b-Kv, although the mean difference was less than 50% and not statistically significant (**Fig. S8D**). These results indicate that somatically targeted ASAP4 GEVIs express similarly well as ASAP3 in neurons *in vivo*. The same trend in brightness was seen under 2-P illumination at 940nm in HEK293a cells (**Fig S7D**), suggesting that ASAP family GEVIs are consistent and usable across both imaging modalities.

To determine how well an ASAP4-family GEVI performs in brain tissue, we compared the performance of ASAP3-Kv and ASAP4b-Kv in mouse hippocampal slices during current-evoked APs. Adeno-associated virus serotype 8 (AAV8) capsids expressing ASAP3-Kv or ASAP4b-Kv in a cre-dependent manner were injected into mouse hippocampus together with AAV-CaMKII-cre. Acute slices were prepared approximately 4 weeks later, then APs were evoked with current pulses of 1-pA amplitude, 1-ms duration, and 10, 20, or 50 Hz frequency during 1-P microscopy. Responses were similar between the two indicators at 10 Hz (**Fig. S8E**), and spikes could be discerned up to 50 Hz (**Fig. S9A**). SNR values were similar between ASAP4b-Kv and ASAP3 across firing rates (**Fig. S9B**). We also compared ASAP4b-Kv and ASAP3-Kv responses under 2-P imaging of patch-clamped neurons in hippocampal slices. Responses to APs were again similar between the two indicators at 10 Hz (**Fig. S8F)** and 50 Hz (**Fig. S9C**), as were SNR values (**Fig. S9D**), and were larger than 35% in amplitude (**Fig. S8F**). When applying current steps to evoke AP trains in place of individual current injections, ASAP4b-Kv again demonstrated similar responses and SNRs as ASAP3-Kv (**Fig. S10**). These results indicate that the performance of ASAP4b-Kv initially characterized in HEK293 cells is well preserved in neurons in brain tissue.

### 1-photon imaging of interneuron activity with ASAP4e-Kv in awake mice

While 1-P voltage imaging with ASAP-family GEVIs in the brain has not been reported previously, their higher molar brightness than opsin-only GEVIs, which have been used for *in vivo* 1-P imaging, suggests they should work as well. ASAP4b-Kv and ASAP4e-Kv have similar per-cell SNRs for spikes, but in 1-P where scattered signals can come from out-of-focus neurons, the lower baseline brightness of ASAP4e-Kv should be preferable. We expressed ASAP3-Kv or ASAP4e-Kv in somatostatin-positive interneurons of mice in the dorsal CA1 region of the hippocampus. Imaging was done through a cranial window using a CMOS camera while mice were awake, head-fixed and running. Illumination by a LED at a power density of 100 mW/mm^2^, an order of magnitude less intense than previously used for 1-P opsin imaging (*29*), easily allowed ASAP imaging with exposure times of 1 ms (**Fig. 2A**), enabling 1-kHz sampling rates on a standard CMOS camera.

**Figure 2.**
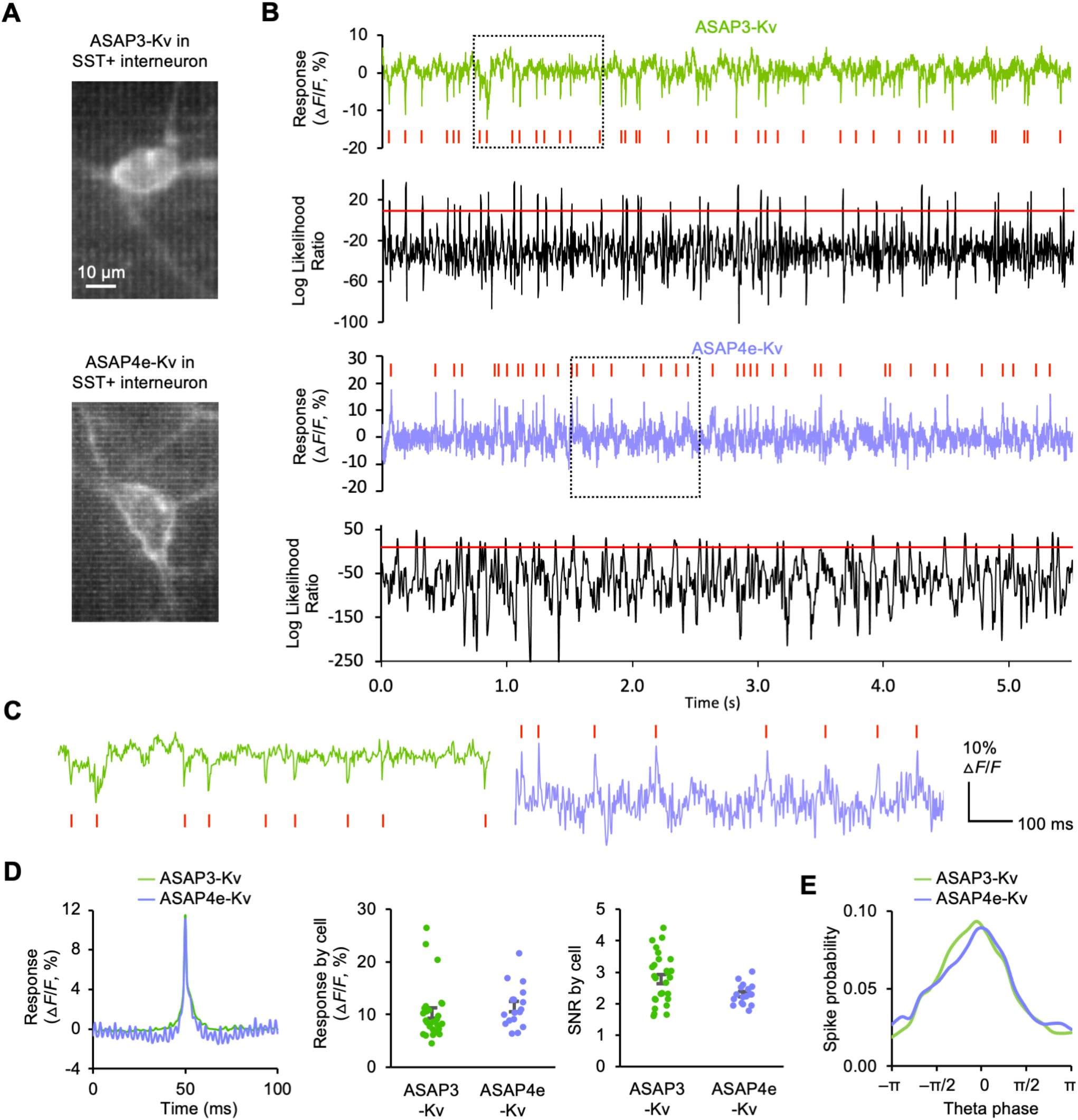
One-photon imaging of ASAP GEVIs in SST+ hippocampal interneurons in vivo. **(A)** Example images of cells expressing ASAP GEVIs imaged through a deep window. **(B)** Example traces recorded at 1 kHz sampling rate then downsampled to 500 Hz but not filtered. Below each trace is the log likelihood ratio probability trace for the 500 Hz downsampled trace in black, with the threshold drawn as a red line. The threshold was chosen such that after breaking the data vector into pieces with lengths equal to the length of the spike template, there was a 1/(20 x number of pieces) chance of getting a false positive in any given piece, which was done to correct for multiple comparisons. Signals were obtained from manually drawn ROIs around the soma. **(C)** Zoomed in examples, corresponding to the segments boxed in with a dotted black line in (B). Traces indicate raw data that has not been filtered in any way. **(D)** Spike metrics from n = 28 cells for ASAP3-Kv, and n = 17 cells for ASAP4e-Kv. For the purposes of making accurate comparisons between indicators without taking into account photobleaching, only data from the first 11 s of each cell was used. Left: The average waveform of detected spikes, with the response direction of ASAP3-Kv flipped for display purposes. Middle: The average response size ΔF/*F*_0_ by cell for both ASAP3-Kv (left) and ASAP4e-Kv (right). The difference was not significant (p = 0.0992 Mann-Whitney test). Right: The average empirically calculated SNR per cell. Note that this was calculated on the raw, unfiltered data after correcting it to 100 mW under Poisson assumptions to account for differences in illumination intensity between cells. **(E)** Histogram of Spike probability over theta phase. Phase was broken into 20 timing bins from 0 to 2π radians in the theta band, defined as 6 to 11 Hz. ASAP3-Kv contributed 10886 spikes from 28 cells, while ASAP4e-Kv contributed 6588 spikes from 17 cells. Both indicators revealed a similar trend for spike preference near the peak at 0 radians.

To detect APs, we employed a template-matching log-likelihood strategy based on the log-likelihood probability work of Wilt et. al., 2013 (**Fig. 2B,C**, see methods), but used Gaussian assumptions about the noise rather than Poisson ones. Briefly, putative APs for training were identified from optical spikes occurring selectively in the negative direction for ASAP3-Kv and in the positive direction for ASAP4e (**Fig. 2B,C**). As deflections in the incorrect direction are due to noise, this allowed the establishment of an amplitude threshold beyond which optical spikes in the correct direction are predominantly reporting APs. At least 10 of the largest of these spikes were chosen to serve as templates for the log-likelihood calculation. New template spikes were chosen for each trial, and if good templates were not available in a trial, then the templates from the same cell but a different trial were used. While this results in conservative detection, we wanted to be sure we did not have many false positives. The log-likelihood function itself exhibited a symmetric distribution with positive outliers; a threshold was set to identify these outliers as AP events (**Fig. 2B,C**). Using this algorithm, optical spike amplitudes (Δ*F*_*event*_/*F*_*rest*_) averaged 10.2 % for AS AP3-Kv and 11.5 % for ASAP4e-Kv, and empirically measured SNR values averaged 2.8 and 2.3, respectively (**Fig. 2D**).

To determine if GEVIs can relate spike generation to subthreshold modulation in the same neurons, we also examined spike onset probability as a function of theta phase. Fluorescence traces were bandpass filtered from 6-11hz, then the Hilbert transform applied to obtain the instantaneous amplitude and phase, and a histogram of the theta phase values when spikes occurred was created. Both ASAP3-Kv and ASAP4e-Kv showed a preference for spiking near the peak of theta, which was set to 0 radians, although spikes can also be seen at other points in theta phase as well (**Fig. 2E**). Note that most studies examining theta have had to look at LFP theta, where spiking preferentially occurs in different phases of the theta oscillation, depending on the cell type (*17*). This extracellular measure of theta will be reversed in polarity relative to the intracellular theta. Intracellular studies examining the relationship between spike probability and theta oscillations in SOM+ interneurons of the hippocampus have found that on average, SOM+ cells prefer to spike at the peak of the theta oscillation (*18*). This makes sense intuitively, because this is when the cell is the most depolarized, and thus the closest to the AP threshold. This is also what we see in our own data in vivo (**Fig. 2E)**. This highlights a great use case for GEVIs in being able to measure these oscillations intracellularly. Future studies using GEVIs for population imaging will be able to see how these intracellular subthreshold dynamics affect networks in vivo during behavior, something extracellular electrode arrays and calcium indicators are unable to do.

ASAP3-Kv and ASAP4e-Kv showed remarkable photostability in vivo, allowing minutes of recording when illuminated at powers between 50 and 247 mW/mm^2^. We also recorded ASAPb-Kv, ASAP4e-Kv and ASAP3-Kv continuously for 30 seconds in Sst+ neurons of the hippocampus (**Fig S11C**). Recordings for ASAP4b-Kv and ASAP4e-Kv were longer, but we cut out the initial photoadaptation effect in traces that showed it to allow them to be aligned to ASAP3-Kv and provide for a direct comparison. All traces in Fig S9C were corrected to show an illumination power of 100mW/mm^2^ under linear assumptions e.g. a 2-fold power increase causes 2-fold faster photobleaching along a monoexponential curve. After 30 seconds, ASAP3-Kv, ASAP4b-Kv and ASAP4e-Kv had bleached to 74.18%, 61.18% and 77.66% of their original brightness on average, respectively (**Fig S11C**). Also consistent with *in cellulo* observations, the photobleaching phase was faster for ASAP4b-Kv than ASAP4e-Kv or ASAP3-Kv, although in all cases the half-decay time when fit to a monoexponential curve was > 1 min at 100 mW/mm^2^ of LED illumination. For real world use case illustrative purposes, we show an example in mice expressing either ASAP4b-Kv or ASAP4e-Kv in Sst+ neurons of the hippocampus during running (**Fig S11A-B**). The ASAP4b-Kv and ASAP4e-Kv examples were exposed to 88 s of illumination (in 8 bouts of 11 s imaging trials during running behavior with 8.1 s darkness gaps in between running bouts). Both traces are shown at 500hz, with ASAP4b-Kv illuminated with 247mW/mm^2^ of power, and ASAP4e-Kv illuminated with 95mW/mm^2^ of power. Traces were not filtered in any way, and were divided by the average of the first 200 ms of the data trace shown. The same numerical average was used to divide the trace shown from 84.2 to 86.2 s, to show the percentage of the original brightness retained. Higher power at 247mW/mm^2^ increased the bleaching rate, but after 86.2 s ASAP4b-Kv still retained 33.04% of its original brightness. ASAP4e-Kv imaged at 95mW/mm^2^ of power retained 44.93% of its original brightness, in these examples, where spikes can still be seen despite the bleaching (**Fig S11A-B**).

### Simultaneous calcium and voltage imaging in vivo with ASAP4e-Kv

Calcium indicators are widely used to reveal neuronal activity *in vivo*, but it is well known that they track calcium concentrations, not voltage. In some instances, calcium and voltage have even been shown to do surprisingly different things (*45*). Additionally, ASAP4’s slower offset kinetics might allow us to get away with really low frame rates, and still obtain information that could not be obtained with a GECI, allowing for its application on standard imaging setups. We thus explored whether an ASAP4 GEVI can be imaged simultaneously with a red GECI using a standard raster-scanning 2-P microscope equipped with a resonant galvo using frame rates from 15-99hz.

We co-expressed the dimmer but more responsive ASAP4e-Kv and the red GECI jRGECO1a in cortical layer 2/3 neurons of mice, then performed 2-P imaging in awake, head-fixed mice. At 1,000 nm excitation, we acquired 128×128-voxel frames while detecting simultaneously emitted green and red fluorescence, corresponding to ASAP4e-Kv and jRGECO1a, respectively. We imaged 13 spontaneously active neurons in one animal, with data from one representative neuron shown in Figure 5. ASAP4e-Kv photostability was excellent; in 100-second continuous recordings, a monoexponential fit showed that 90.86% of beginning fluorescence was retained (**Fig. 3A, Movie S1**). Despite the 1,000 nm excitation being non-optimal for ASAP, we observed that each GECI transient was temporally preceded by an ASAP4e-Kv transient (**Fig. 3A, Movie S1**). There was a clear relationship between the number of closely spaced spikes and the amplitude of the GECI transient, such that smaller voltage transients were associated with only very small increases in GECI signals, and sometimes almost none at all (**Fig. 3B**).

**Figure 3.**
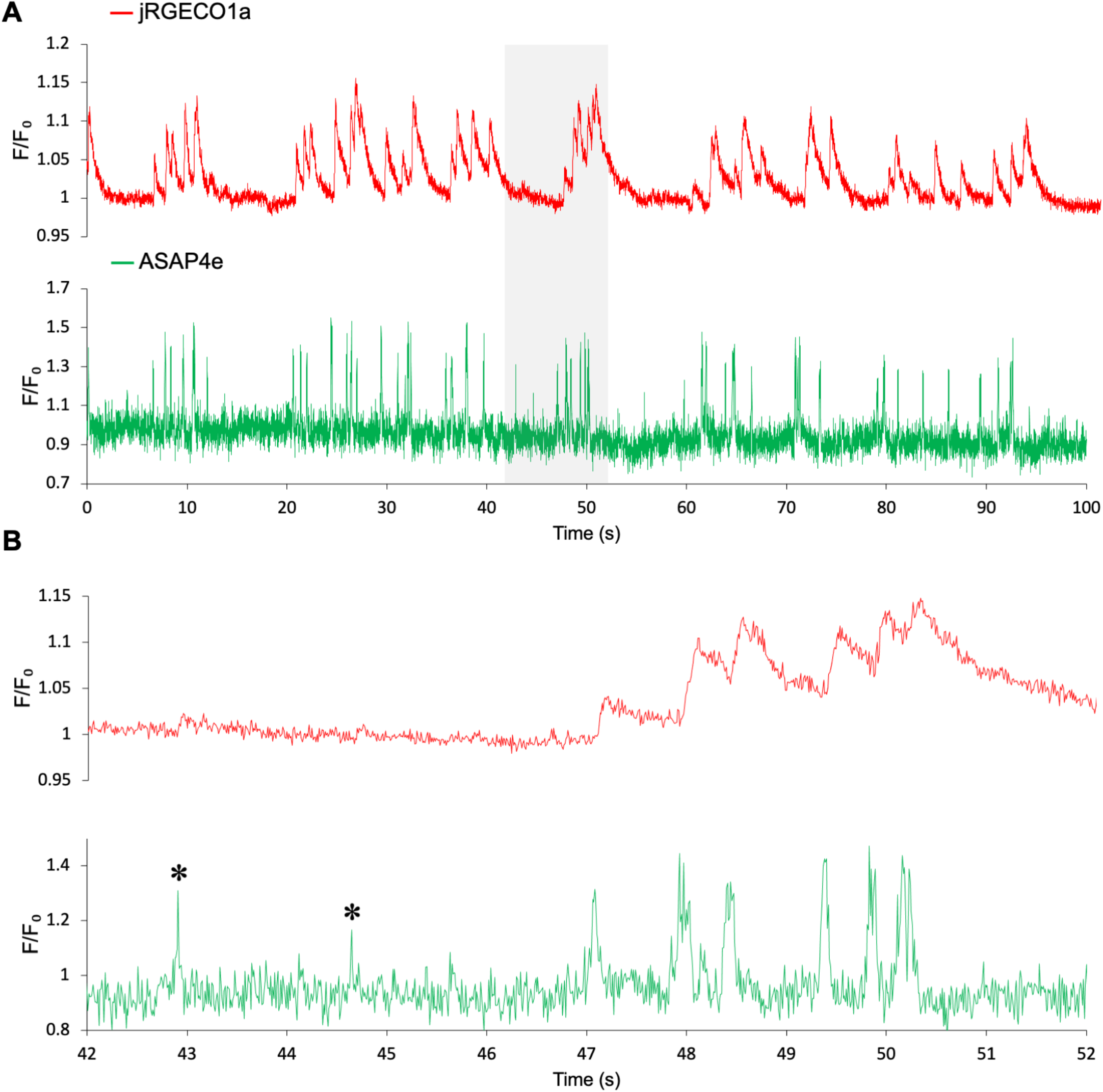
Simultaneous voltage and calcium imaging in cortical pyramidal neurons in vivo. (A) ASAP4e-Kv and jRGECO1a were co-expressed in layer-2 cortical neurons of mice. Imaging was done through a cranial window over V1 with a standard galvonometer-based scanner at 99 fps using a wavelength of 1000 nm to excite both ASAP4e-Kv and jRGECO1a simultaneously, despite 1000nm being suboptimal for ASAP4e. Traces shown have been normalized to 1 by dividing by the average of seconds 4-5, but are otherwise unmodified and uncorrected for photobleaching so that photobleaching may be observed, and represent raw and unfiltered data. The gray shaded area represents the zoomed in portion in panel B. (B) The asterisks (*) mark voltage trace events that show little to no response in the calcium (jRGECO1a) channel. To the right of that is shown a burst of activity, where ASAP4e-Kv is able to track electrical events much faster than jRGECO1a, and always precedes them temporally. Link to this dataset as Movie S1: https://drive.google.com/drive/folders/1jlHcrM0PZVaVMfJpqQuBJOHBNvu6ZvFw?usp=sharing

### Imaging place cell population dynamics with ASAP4b-Kv

Finally, we used ASAP4 and jRGECO1b to simultaneously record voltage and calcium activity in CA1 pyramidal neurons as mice navigated through a virtual reality environment to find hidden rewards (**Fig. 4A,B**). For this task, we chose ASAP4b-Kv as we desired to scan as many neurons as possible, and the higher molecular brightness at rest may be more useful for detecting multiple neurons over a large region, similar to the use case for jGCaMP7b (*11*). With minimal modifications to a commercially available resonant-scanning 2-P microscope (Neurolabware), we were able to continuously image rectangular fields of view containing >50 cells at 989 Hz for tens of minutes (**Fig. 4C**). To minimize power delivery and to use a commonly available imaging setup, we used one excitation laser at 940 nm at a minimum power capable of detecting ASAP4b-Kv at baseline, and split green (ASAP4b-Kv) and red (jRGECO1b) signals to separate detectors.

**Figure 4.**
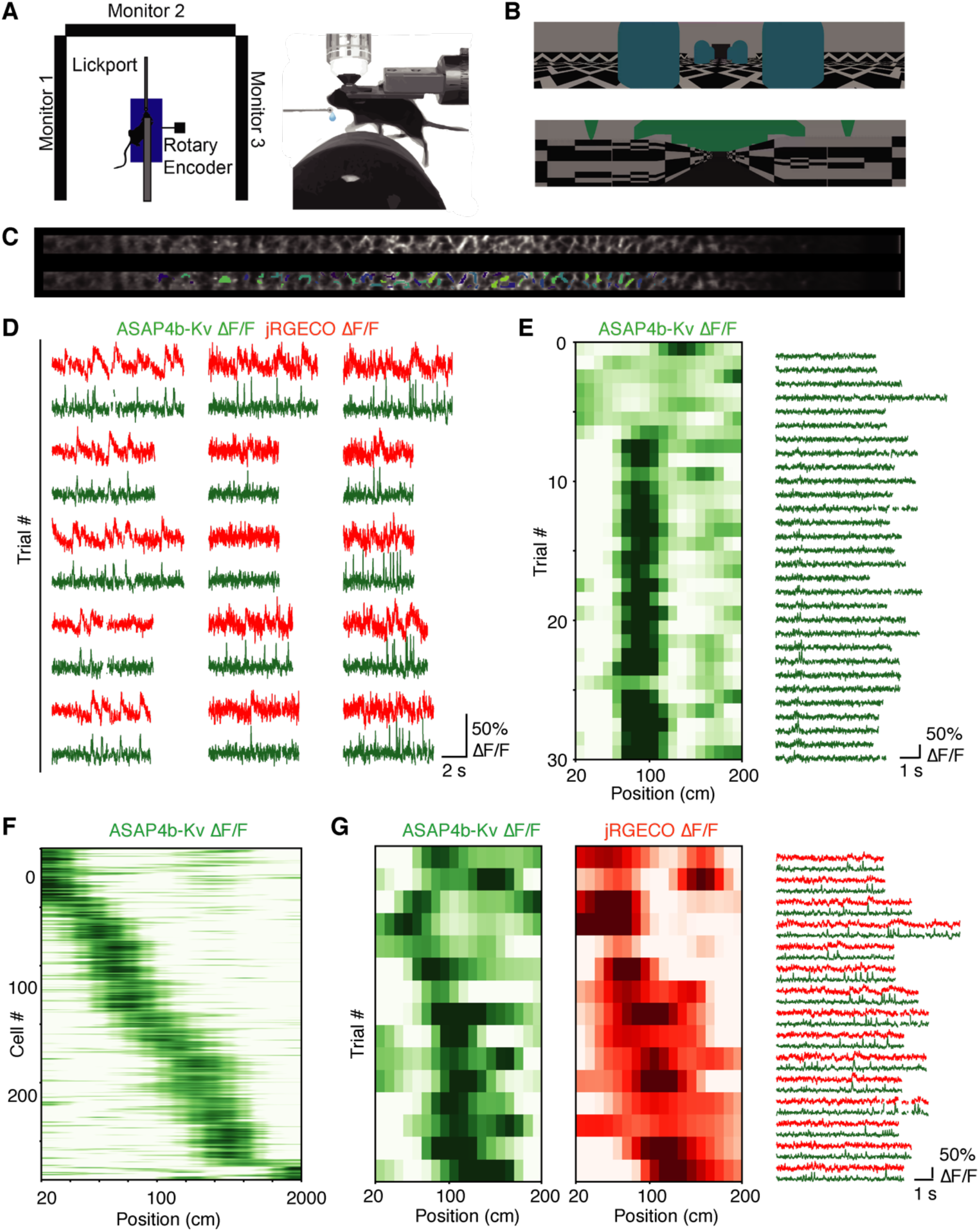
Precision mapping of hippocampal place cells by voltage imaging. **(A)** Virtual reality system. A head-fixed mouse is allowed to run on a fixed axis cylinder. A rotary encoder translates cylinder rotation to movement through a virtual linear track displayed on three screens surrounding the mouse. **(B)** The mouse’s view at the beginning of the two virtual linear tracks used for data collection. Data was pooled across both virtual environments. **(C)** Example ASAP image without (top) and with (bottom) ROIs highlighted. **(D)** Example ROIs displaying high SNR in simultaneously recorded ASAP4b-Kv and jRGECO1b channels. Δ*F*/*F*_0_ traces from a subset of trials in a single session. ASAP4b-Kv and jRGECO1b display highly correlated transients but with different kinetics. Missing data is from frames estimated to be corrupted by brain motion. **(E)** Example place-selective ROI from the FOV shown in (C). Left, heatmap indicates ASAP4 Δ*F*/*F*_0_ averaged within each spatial bin on each trial. Each row is the position-binned activity for that cell on a single trial. Right, Δ*F*/*F*_0_ traces corresponding to these trials. Traces are of different durations due to brain motion. **(F)** Trial-averaged (bootstrapped) ASAP4b-Kv activity of every place-selective ROI plotted as a function of position (n = 283 ROIs) recorded across all mice (n = 3) and all imaging sessions. Each row indicates the activity of a single ROI. ROIs are sorted by their location of peak activity and z-scored. **(G)** Left, place fields from a cell where similar patterns of activity can be seen in both ASAP4b-Kv and jRGECO1b channels with place preferences. Right, Δ*F*/*F*_0_ traces corresponding to these trials. Traces are of different durations due to brain motion.

Applying an analysis pipeline previously developed for calcium imaging to ASAP4b-Kv fluorescence (see Methods), we identified a large population of place cells (∼30% of cells with significant spatial information). As expected from calcium imaging and extracellular voltage recordings, ASAP4b-Kv place cells tiled the environment with their place fields (**Fig. 4D-G**). Calcium and voltage activity rate maps were largely consistent in cells with strong co-expression, but ASAP4b-Kv signals clearly demonstrated higher temporal resolution and better ability to discern closely spaced spikes (**Fig. 4D,G**). Thus, using a standard commercial 2-P microscopy system designed for imaging calcium indicators, we were able to record populations of place cells in the hippocampus by imaging ASAP4b-Kv. Additionally, the signals were large and robust enough that a standard calcium imaging data processing pipeline using Suite2P coupled with a common spatial information metric (*43*), was able to pull out place cell tiling in the green ASAP4b-Kv channel that matched the red jRGECO1b channel (**Fig. 4G**).

## DISCUSSION

In this study, we introduce two responsive, positively tuned GEVIs, ASAP4b and ASAP4e, and use them to detect spiking activity in single trials in vivo by 1-photon and 2-photon imaging. We demonstrate that both ASAP4e and the negatively tuned ASAP3 can be used to detect spikes in multiple hippocampal interneurons by 1-photon imaging for over a minute, establishing the ability to perform voltage imaging with epifluorescence equipment commonly used for imaging GFP-based GECIs. Importantly, the size of the field of view was not limited by the need to focus the illumination source into a small area, but instead by our camera which needed to scan a small FOV in order to operate at 989hz. Presumably with a better camera, the same illumination could be used, only a much larger FOV containing many more cells imaged, opening the door to large scale population imaging under 1-P illumination with GEVIs.

We also demonstrate simultaneous single-cell voltage and calcium imaging by two-photon microscopy of ASAP4e and ASAP4b with a red GECI. Our results reveal that calcium transients in layer 2 pyramidal neurons of the cortex do not always capture the underlying voltage activity, and that ASAP4 can be used simultaneously to image voltage on a standard galvanometric 2-P imaging setup, even at frame rates as low as 99hz. While temporal resolution will obviously be affected at low frame rates, we believe that this will allow for calcium and voltage to be studied simultaneously in all cell types, and we anticipate some surprising findings resulting from that in the future. We also find that voltage imaging can be used to map place fields of hippocampal pyramidal neurons using a standard calcium indicator processing pipeline relying on spatial information (*43*). This provides the voltage imaging equivalent to the spike-based place field mapping performed by extracellular electrodes and GECIs.

With regard to the relative coverage of GEVI and GECI imaging, it is informative to compare molecular performance and identify inherent limiting factors. In response to a single 2-ms AP, ASAP4e responds with a relative increase of 25% at peak. This is similar to the response relative to baseline of GCaMP6f, which is also ∼25% for a single AP (*9*). However, in terms of per-molecule brightness and kinetics, ASAP4e and GCaMP6f differ in opposite ways. To produce its 25% response, ASAP4e fluorescence increases from 20% to 25% of the maximum brightness of its GFP fluorophore. In contrast, GCaMP6f increases from ∼2% to ∼2.5%. Thus the change in photon flux per molecule in response to an AP is an order of magnitude larger for ASAP4e. However, an opposite relationship is observed in the persistence of the response; GCaMP6f is an order of magnitude slower in its signal decay compared to ASAP4e (off time-constants of 150 ms vs. 15 ms at room temperature). Another major difference between GEVIs and GECIs is their abundance in a neuronal soma; it has been estimated there are ∼20 times more GECI molecules (*6*). Thus the higher molecular performance of GEVIs is in practice balanced by the shorter imaging intervals that are needed to capture spiking activity with GEVIs. GEVI imaging is then further disadvantaged by an inherently lower molecular abundance than GECIs.

Nevertheless, it is still possible that future improvements to ASAP-family GEVIs could help reduce the 2-P coverage disadvantage by producing larger per-AP responses, as this would reduce dwell time requirements while maintaining SNR. In response to steady-state voltages from –70 to +30 mV, ASAP4 operates from 20% to 57% of the maximum brightness of its GFP fluorophore. Sharpening the F-V curve to further decrease basal fluorescence at –70 mV and increase fluorescence at +30 mV will produce larger responses to voltage changes. Accelerating activation kinetics will allow for more of the already substantial steady-state fluorescence range (178% for ASAP4e) to be sampled during APs. Finally, selectively slowing deactivation kinetics further would allow for more widely spaced imaging intervals, although at a cost in resolving closely spaced spikes. Such optimizations would also aid 1-photon imaging, improving the SNR for spike detection or allowing lower illumination powers, thereby increasing reliability of event detection or duration of recording.

All of the imaging systems used in this study were standard systems routinely used for calcium imaging, many of which have been around for over a decade. We believe that ASAP4 now allows for any lab currently imaging GECIs to plug ASAP4 into their pipeline, with minimal equipment optimization needed. However, if the interest is to maximize the number of cells imaged per session with optimized SNR, then more sophisticated imaging methods, such as patterned or random-access illumination, would allow for even higher coverage.

In summary, we perform voltage imaging using ASAP4-family GEVIs on existing 1-P and 2-P equipment previously developed for imaging GECIs, demonstrating that voltage imaging is within the technical abilities of a large number of neuroscience laboratories. GEVI imaging should be widely applicable for recording neuronal spiking activity with higher temporal precision than possible with calcium imaging. Future work will aim to improve genetically encoded voltage indicators in detection reliability, to allow for larger populations of cells to be imaged while maintaining high SNR.

### Supplementary Note 1

We wanted to shift the F-V curve leftwards so that V_1/2_ would lie within the physiological range. In energetic terms, this is equivalent to stabilizing the deprotonated (bright) state of the chromophore. Position T206 is known to interact with the chromophore and alter its pKa, and so we chose it for saturated mutagenesis in our screening system, hoping to find an amino acid that further stabilized the deprotonated state of the chromophore. We were rewarded with a T206H mutation that resulted in a dramatically left-shifted GEVI, which we designated ASAP4.1. *F*_+30_ reached 91% of F_max_, and ΔF_100_ improved to 35% of *F*_max_ (**Supp Table 1)**. We still had not regained the dynamic range of ASAP4.0, although we had increased the brightness significantly (**Supp Table 1**).

To search for further improvements, especially to dynamic range, we next mutagenized Phe-148 and Asn-149, the first two GFP-derived residues after the S3-cpGFP junction in the linker region. These sites are directly adjacent to the sites that we screened to initially flip the response, and we reasoned that in addition to interacting with them, Phe-148 and Asn-149 should be critical in translating VSD movement into chromophore stability and therefore improving our indicator. After screening all 400 combinations of possible amino acids by electroporation, we identified a F148P N149V mutant (ASAP4.2) as exhibiting a preferential decrease in fluorescence at –70 mV (*F*_–70_) compared to fluorescence at +30 mV (F_+30_). This resulted in a sharper *F-V* curve and a larger Δ*F*_100_.

To reduce *F*_–70_ even more, and thereby further improve Δ*F*_100_, we turned to mutating the voltage-sensing domain (VSD) of ASAP4.2. We hypothesized that if we could stabilize the down conformation of the S4 helix at – 70 mV, we could make it dimmer there.

Modeling ASAP with MODELER suggested position 413 as being very important for creating an energy barrier to VSD movement. It was also previously reported to modulate the voltage input tuning of the Ciona voltage-sensing domain, which is structurally similar to our own (*15*). This position needs to move from the cytosol into the membrane bilayer during upward S4 helix movement, and interacts with the external environment. This suggests that hydrophobic/hydrophilic amino acids should have large effects on VSD movement at this position, which should translate into changes in fluorescence. Patch-clamp electrophysiology of all 20 amino acids at position 413 revealed two mutants with the desired dimming at –70 mV and increased Δ*F*_100_, F413V (ASAP4.3) and F413G (ASAP4b). F413M (ASAP4e) was also interesting, as it dramatically reduced brightness at both –70 and +30 mV, although more at +30 mV. This had the undesirable effect of reducing the Δ*F*_100_, but the relative fluorescence change (Δ*F*_100_/*F*_–70_) was improved.

Position 401 was also suggested to influence voltage tuning by modulating the hydrophobic plug there (*21*). Since both could be working together to control and tune the ability of the S4 helix to transition from the cytosol to the lipid bilayer of the cell, we screened all possible combinations of positions L401 and F413 by electroporation in our screening system. To our unpleasant surprise, no combinations gave a better response than ASAP4.3, ASAP4b and ASAP4e, which only had the single 413 mutations of F413V, F413G and F413M, respectively. Still seeking further improvements, we also screened all 400 combinations of position 65 and 69 in the beta barrel, but all resulting mutants were so dim that we did not pursue them any further.

**Table S1.**
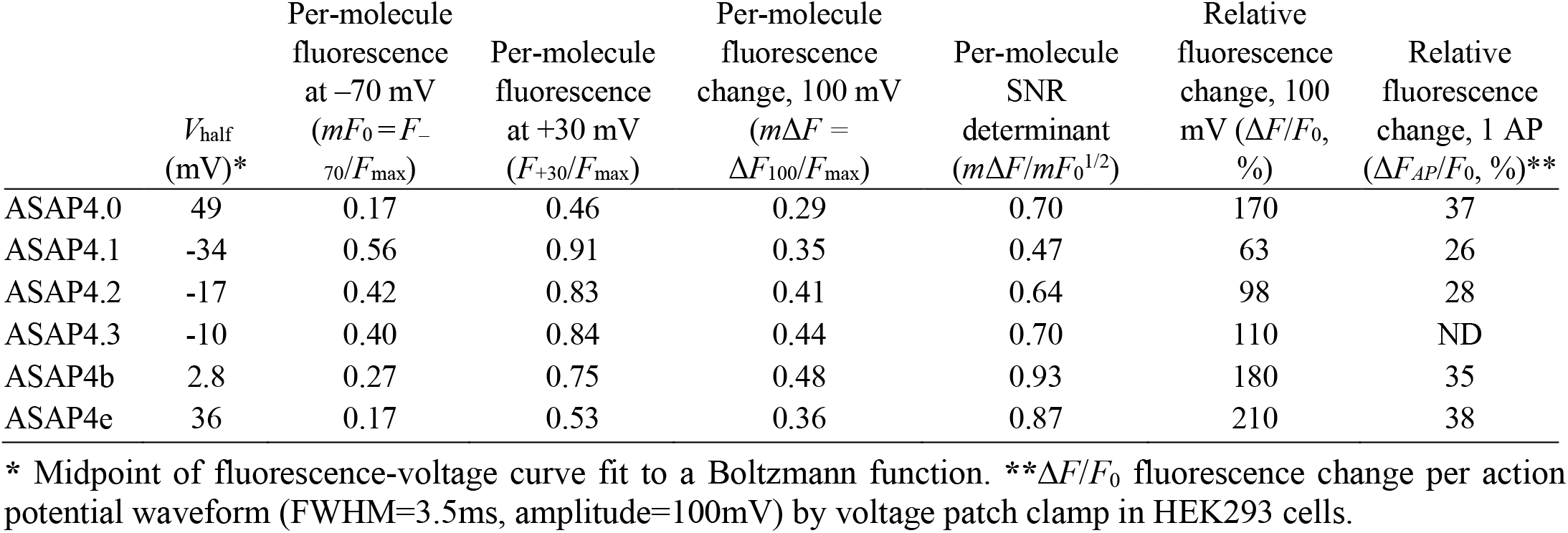
Relative fluorescence per molecule of ASAP4 variants.

**Table S2.** Kinetics for ASAP4 variants.

|  | ASAP4.2<br>at 22 °C | ASAP4.3<br>at 22 °C | ASAP4b<br>at 22 °C | ASAP4e<br>at 22 °C | ASAP4b<br>at 37 °C | ASAP4e<br>at 37 °C |
| --- | --- | --- | --- | --- | --- | --- |
| Depolarization (−70 to 30 mV) |  |  |  |  |  |  |
| $\tau_{\text{fast}}$ (ms) | 2.24 ± 0.18 | 2.89 ± 0.14 | 2.62 ± 0.15 | 3.93 ± 0.27 | 1.53 ± 0.05 | 1.49 ± 0.03 |
| $\tau_{\text{slow}}$ (ms) | 12.95 ± 0.42 | 17.74 ± 0.40 | 21.21 ± 0.83 | 20.64 ± 0.68 | 6.13 ± 0.06 | 5.07 ± 0.03 |
| % fast | 21.80 ± 1.36 | 14.93 ± 0.64 | 19.12 ± 1.64 | 14.05 ± 0.51 | 12.48 ± 0.69 | 14.47 ± 0.57 |
| Repolarization (30 to −70 mV) |  |  |  |  |  |  |
| $\tau_{\text{fast}}$ (ms) | 1.00 ± 0.36 | 5.26 ± 1.13 | 5.69 ± 0.76 | 8.40 ± 0.68 | 1.54 ± 0.10 | 3.66 ± 0.06 |
| $\tau_{\text{slow}}$ (ms) | 15.73 ± 0.42 | 17.45 ± 0.53 | 24.50 ± 0.60 | 17.47 ± 0.31 | 7.38 ± 0.07 | 7.01 ± 0.33 |
| % fast | 31.30 ± 10.21 | 27.46 ± 4.30 | 8.76 ± 1.28 | 20.60 ± 3.93 | 6.52 ± 0.68 | 68.60 ± 2.73 |
| Hyperpolarization (−70 to −100 mV) |  |  |  |  |  |  |
| $\tau_{\text{fast}}$ (ms) | 0.35 ± 0.19 | 3.02 ± 1.40 | 3.83 ± 0.45 | 0.49 ± 0.20 | 1.63 ± 0.07 | 3.69 ± 0.02 |
| $\tau_{\text{slow}}$ (ms) | 18.15 ± 8.54 | 13.60 ± 1.64 | 18.32 ± 0.83 | 13.65 ± 0.34 | 6.80 ± 0.20 | - |
| % fast | 44.58 ± 8.90 | 58.09 ± 9.28 | 37.16 ± 2.71 | 55.35 ± 11.61 | 27.56 ± 1.54 | 100 |
| Repolarization(−100 to −70 mV) |  |  |  |  |  |  |
| $\tau_{\text{fast}}$ (ms) | 2.91 ± 1.10 | 0.94 ± 0.36 | 3.41 ± 0.37 | 0.58 ± 0.28 | 3.88 ± 0.03 | 2.13 ± 0.03 |
| $\tau_{\text{slow}}$ (ms) | 15.99 ± 4.76 | 12.90 ± 0.81 | 21.01 ± 1.30 | 13.43 ± 1.05 | - | - |
| % fast | 69.43 ± 7.11 | 54.70 ± 9.26 | 46.25 ± 2.36 | 58.29 ± 5.94 | 100 | 100 |

Voltage patch clamp was done at either room temperature (22 °C) or 37 °C with a sampling rate of 2500 Hz. Responses had reached the steady state of either depolarization to +30mV or hyperpolarization to -100mV from -70mV, before voltage was returned to baseline for the measurement of repolarization kinetics. All ASAP4 variants were measured in 7 to 11 HEK293 cells. Values are mean ± SEMs.

**Table S3.**
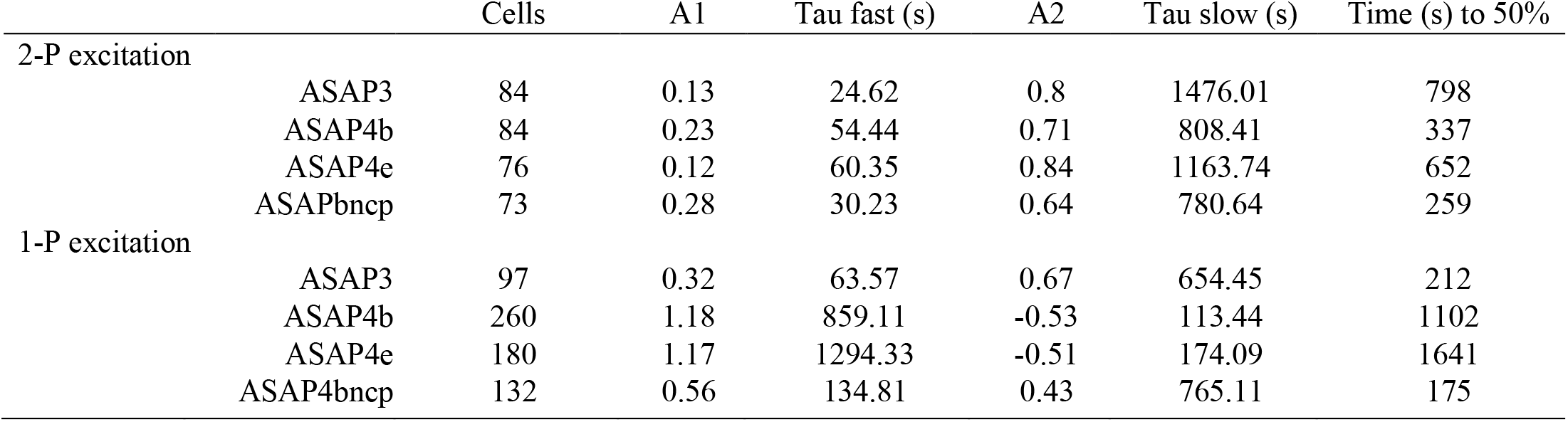
Photobleaching kinetics.

**Movie S1. In Vivo Imaging of ASAP4e-kv and jRGECO1a in Mouse V1**.

https://drive.google.com/drive/folders/1jlHcrM0PZVaVMfJpqQuBJOHBNvu6ZvFw?usp=sharing

**Figure S1.**
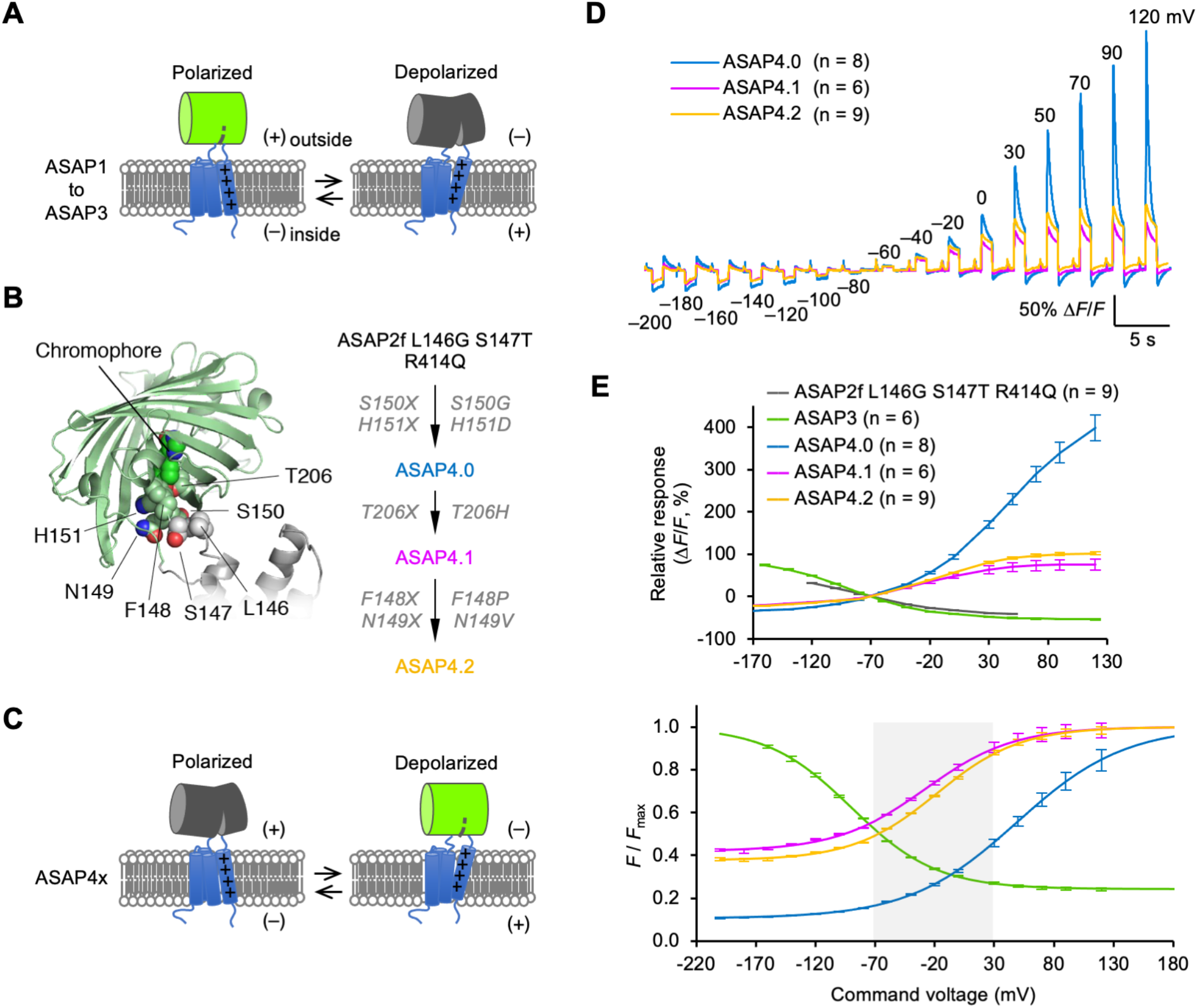
Discovery of a positively-tuned VSD-based GEVI. **(A)** Cartoon of ASAP1 to ASAP3 mechanism, showing the opening of a gap by the chromophore, destabilizing the bright state, when the VSD is in the depolarized confirmation. This gap closes more with membrane hyperpolarization, increasing chromophore bright state stability. **(B)** Left: Schematic showing the beta barrel, chromophore, VSD linkers and the locations of mutated amino acids. Right: A flow chart listing the sites screened, and the resulting ASAP4 mutant from each round of site screens, broken up by arrows. **(C)** The same as (A), but for ASAP4. Now the gap is closed when the VSD is in the depolarized conformation, stabilizing the chromophore’s bright state and increasing indicator brightness with depolarizing voltage across the membrane. **(D)** Steady-state responses from command voltage patch clamping in HEK 293a cells for ASAP4.0 to ASAP4.2. The curves have been normalized to - 70mV. The test pulses before each step are 2ms square steps from –70 to +30mV (E) Top: Δ*F*/*F*_0_ over voltage steps for the parent construct ASAP2f L146G S147T R414Q, as well as ASAP3 and the first three ASAP4 variants. Error bars are SEM. Bottom: The same plot, only displayed as a percent of the maximum brightness of the indicator. Data is displayed as the mean ± SEM. The transparent light gray highlights the physiological range from –70 to +30mV

**Figure S2.**
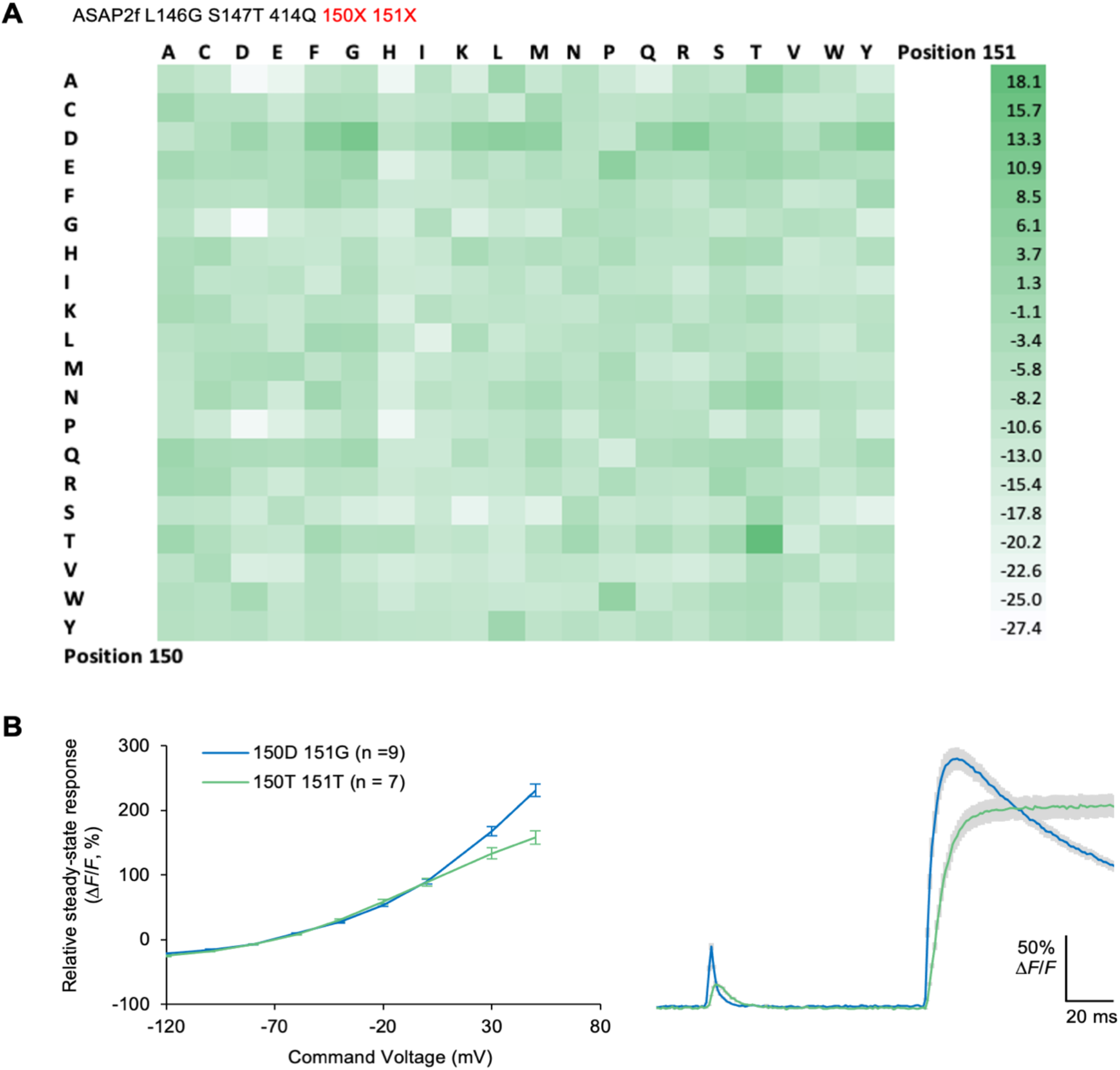
Reversing the Fluorescence Response. **(A)** Beginning with ASAP2f L146G S147T 414Q, a mutant that was also a precursor to ASAP3, we screened 2 amino acids at the interface of the VSD linker and the beta barrel, positions 150 and 151 in ASAP numbering, seeking to open the barrel when the VSD was in the active state. Some of the mutants from this screen gave negative responses like ASAP3, others gave positive-going responses. We chose 150D 151G, as many of the 150D 151x mutants flipped the response, and 150D 151G was the largest of them. While mutant 150T 151T had the largest response overall, a follow up screen showed it to only have a response of 7% ± 3.2 SEM, while DG was still the best. Combined with the slow kinetics of mutant 150T 151T, we chose 150D 151G as the basis for ASAP4.0. Each mutant was repeated in at least 3 wells, as sometimes wells don’t give good responses due to cell health, transfections issues, and other sources of biological variability. **(B)** Left, FV curves for mutant 150D 151G (ASAP4.0), and mutant 150T 151T, normalized to –70 mV. Upon patch clamping, ASAP4.0 outperformed mutant 150T 151T in terms of responsivity (left), and kinetics (right).

**Figure S3.**
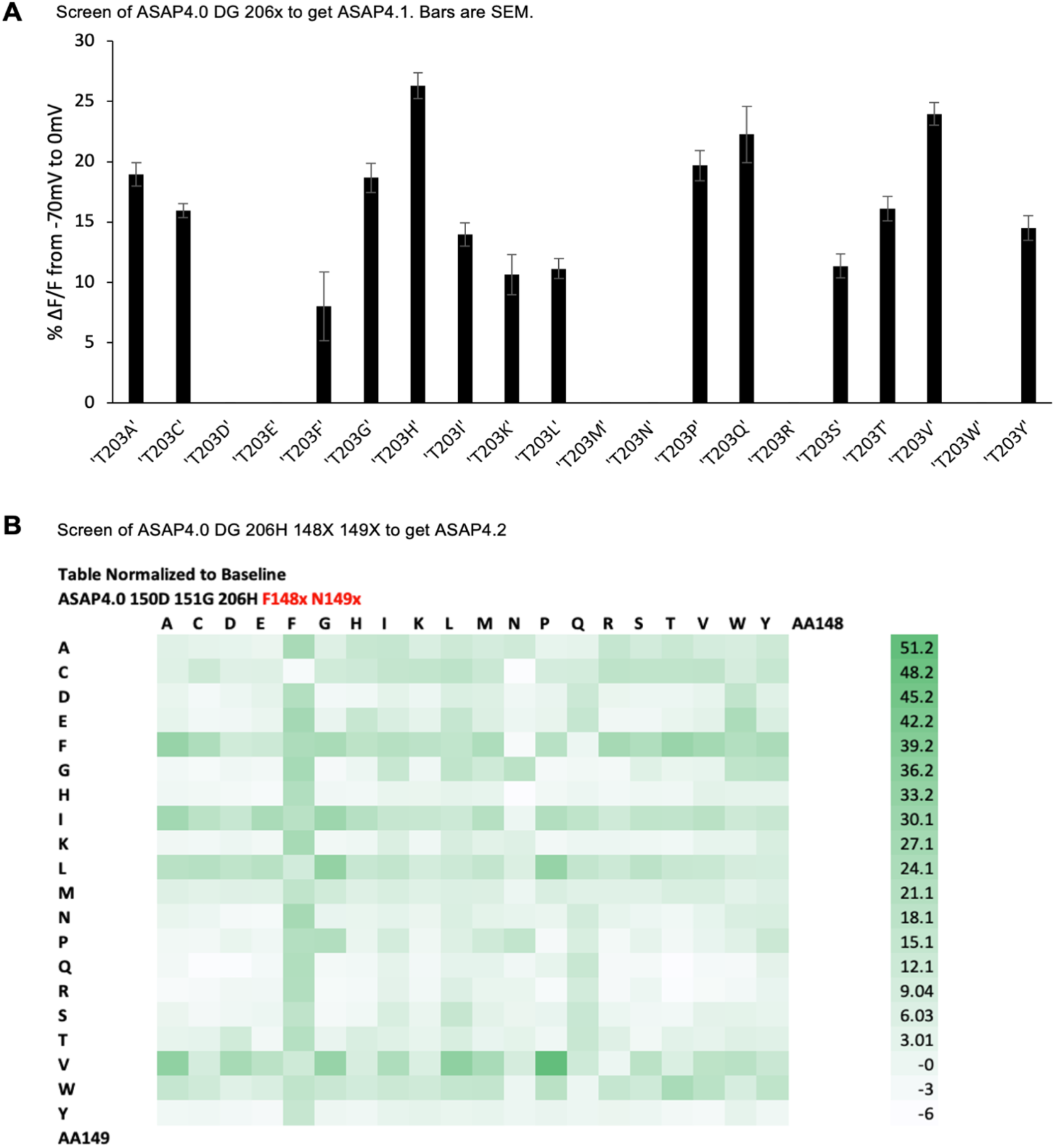
ASAP4.0 to ASAP4.2. **(A)** Position 203T in the beta barrel, which is known to interact with the chromophore, was chosen as a saturated mutagenesis site in an attempt to recover some of the brightness lost when ASAP4.0 was developed. Not only was T203H much brighter, it also gave the largest response when screened. We called ASAP4.0 T203H ASAP4.1 **(B)** Screening all combinations of F148x N149x yielded F148P N149V, creating ASAP4.2.

**Figure S4.**
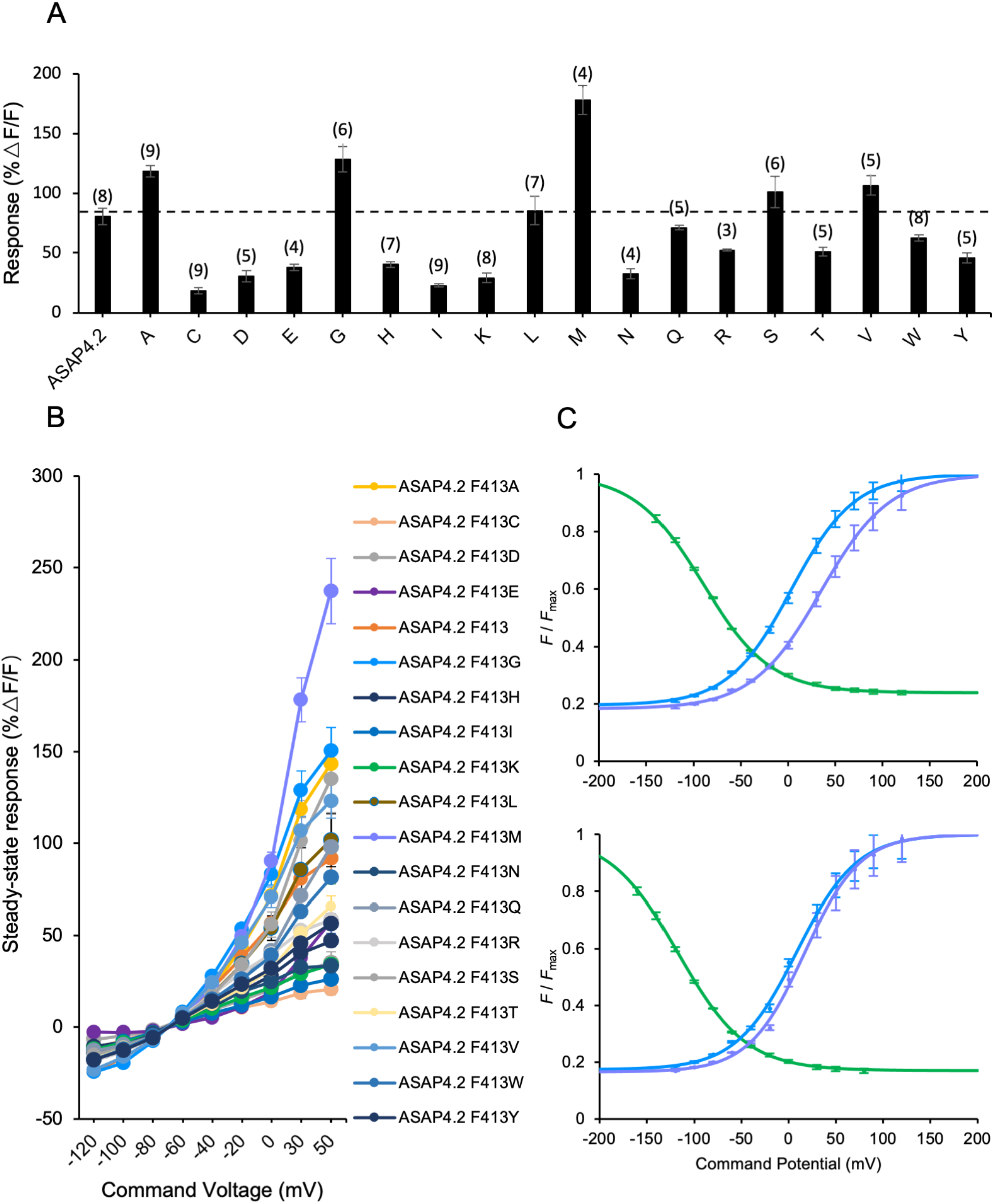
Improvements to ASAP4.2. **(A)** Mean maximum response of ASAP4.2 F413X variants to 100 mV steps in HEK293A cells from a holding potential of −70 mV to +30mV. X-axis labels are the amino acids substituted in the place of F413 in ASAP4.2. The variant with proline didn’t have good expression and thus was not tested. Error bars are mean ± SEM. The number of cells tested is shown in parentheses above the bars. F413A/G/L/M/S/V all gave bigger responses than the control ASAP4.2. **(B)** Mean fluorescent responses of ASAP4.2 F413X variants to a series of voltage steps from -120 to +50, from a holding potential of –70 mV in voltage-clamped HEK293A cells. Error bars are mean ± standard error of the mean (SEM). Only Proline was excluded due to expression issues. The top 3 performers, F413A/G/M, were designated ASASP4.3, 4.4 (4b) and 4.5 (4e), respectively. (C) Top: *F* / *F*_max_ for ASAP3 (green, n = 6 cells), ASAP4b (blue, n = 5 cells), and ASAP4e (purple, n = 5) at 23 °C. Bottom: *F* / *F*_max_ for ASAP3 (green, n = 8 cells), ASAP4b (blue, n = 6 cells), and ASAP4e (purple, n = 6) at 37 °C.

**Figure S5.**
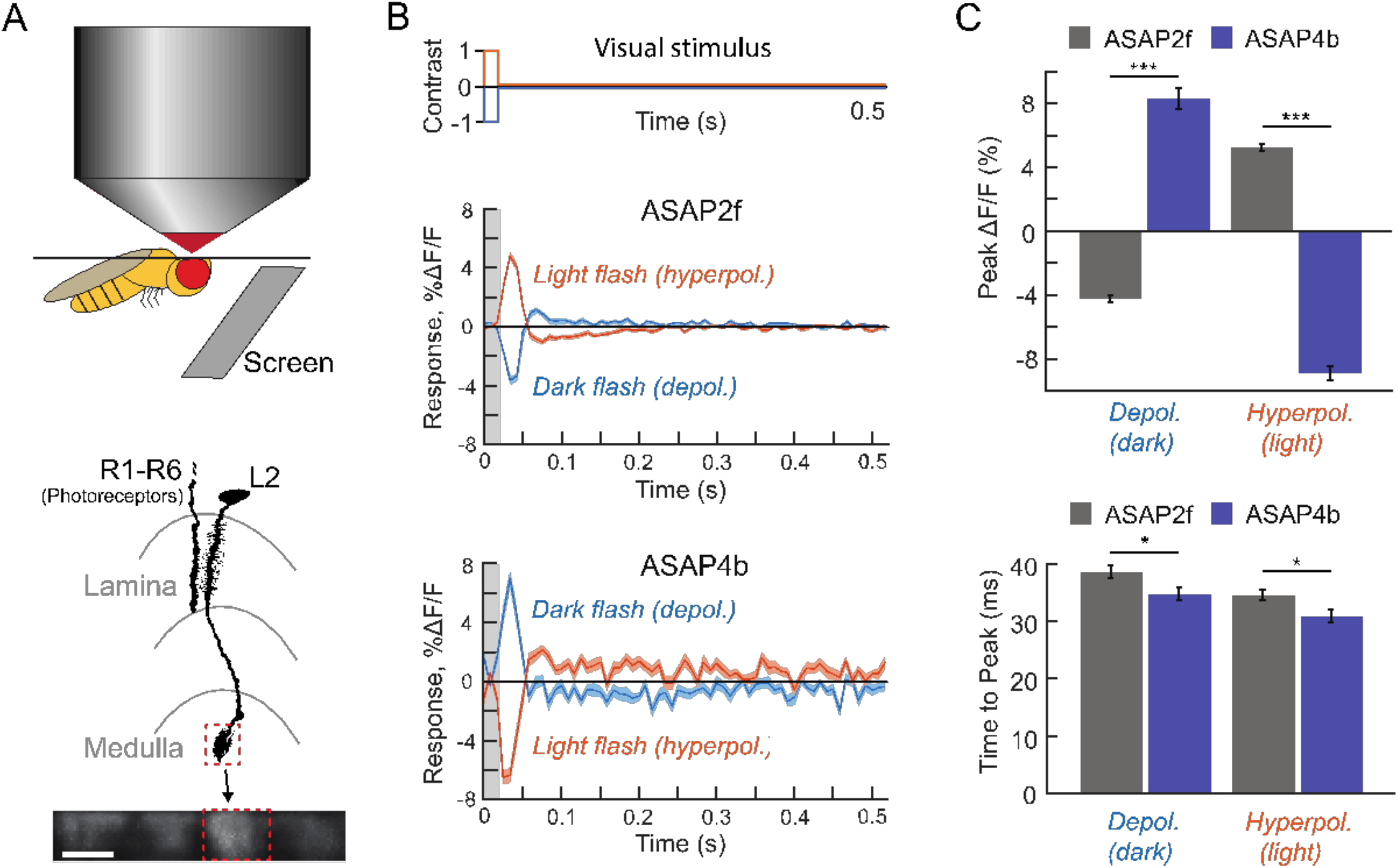
Imaging ASAP4b and ASAP2f in Awake Flies During Visual Stimulation. **(A)** We performed two-photon imaging of visually evoked responses in Drosophila using a flickering visual stimulus on a gray background (above), measuring changes in fluorescence in the non-spiking interneuron L2 (below), which is directly postsynaptic to photoreceptors (R1-6). Bottom: The schematic shows one photoreceptor and L2 cell spanning the lamina and the medulla, two ganglia within the visual system. Both the lamina and medulla are comprised of an array of columns, each containing a single L2 neuron, tiling visual space. We imaged the axonal projections of L2 in the medulla; the image includes the terminal arbors of four neighboring L2 cells. L2 cells depolarize to dark flashes and hyperpolarize to light flashes (*27*). **(B)** Average stimulus-evoked voltage responses, measured in L2. Top: Schematic of the stimulus, displaying contrast over time. ASAP2f (middle, n = 43 cells from 5 flies) and ASAP4b (bottom, 45 cells from 4 flies), show that ASAP4b reports depolarizations with an inverted deltaF/F relative to ASAP2f. That is, ASAP4b becomes brighter when the cell is depolarized and dimmer when the cell is hyperpolarized. We presented a repeated sequence of 20 ms light and dark flashes starting from a mean gray background, indicated with light gray shading near time = 0 in the middle and bottom panels. Blue traces show L2’s response to dark flashes; orange traces show L2’s response to light flashes. The colored line is the average of each cell’s stimulus-aligned average; shaded area is the SEM. **(C)** Top: Peak response amplitudes show that ASAP4b has a significantly larger fluorescence response change than ASAP2f to the same stimulus, and responds in the opposite direction. These metrics are taken from the same single-cell stimulus-aligned data that went into B). The plots show the average and SEM of those values. *** p<0.001 for depolarization (dark) and for hyperpolarization (light), using a two-sample t-test with Bonferroni correction. Bottom: Time-to-peak measurements show that ASAP4b produces responses with slightly slower kinetics as ASAP2f, only with a much larger peak as seen in (B) and (C), as expected from their respective kinetics. These metrics are taken from the same single-cell stimulus-aligned averages that went into B). The plots show the average and SEM of those values. * p = 0.0406 for depolarization (dark) and 0.0268 for hyperpolarization (light), by a two-sided t-test with Bonferroni correction.

**Figure S6.**
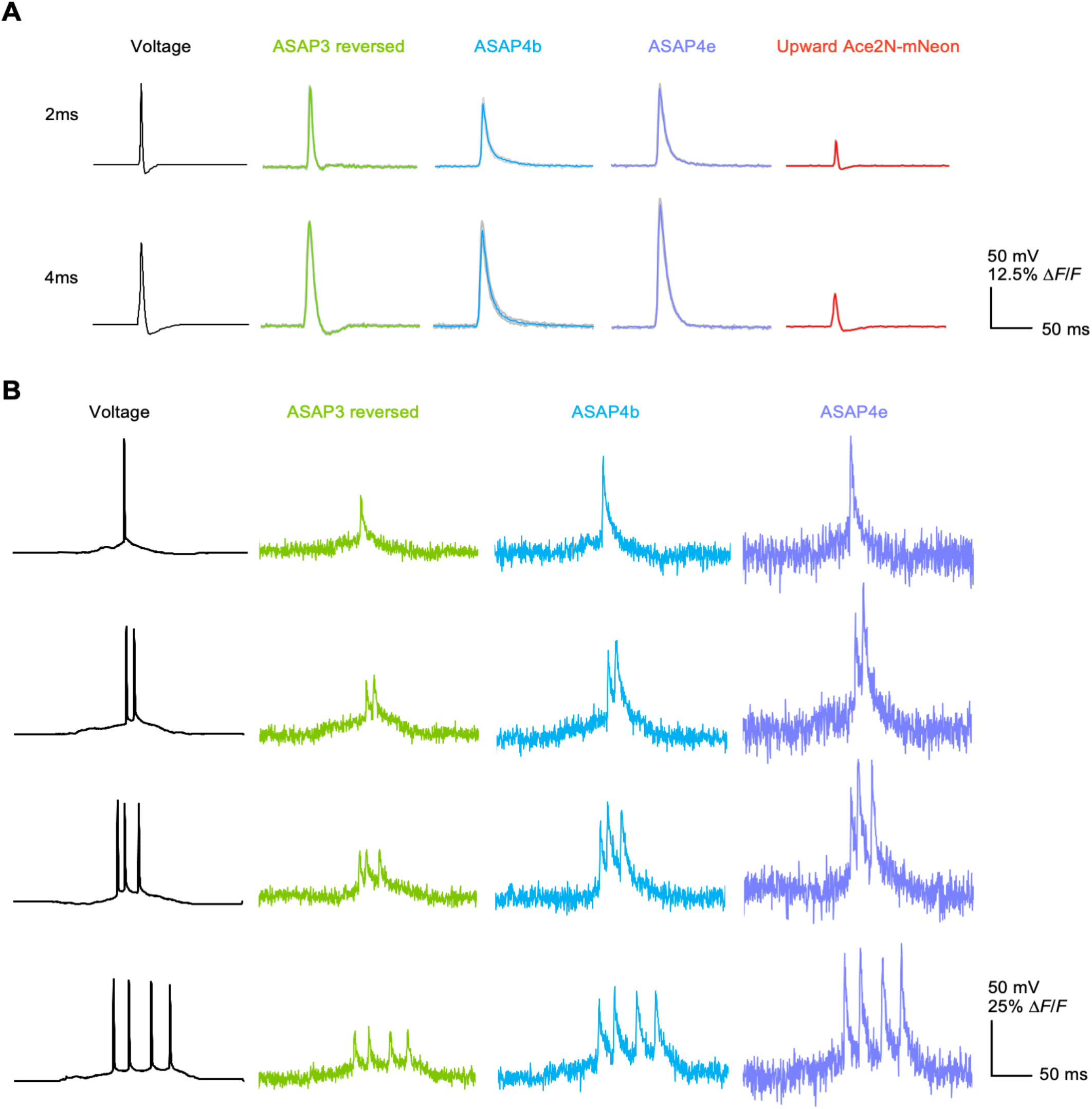
Responses of ASAP family GEVIs to AP waveforms in HEK293A cells. **(A)** Responses of indicators in HEK293a cells to an AP waveform (far left) under voltage clamp that has been modified to have either a 2ms FWHM (top), or a 4ms FWHM (bottom), and ranges from -70mV to +30mV. Each cell had the waveform applied 5 times, with n = 6 cells for ASAP3, n = 7 cells for ASAP4b, n = 5 for ASAP4e, and n = 7 cells for Upward Ace2mNeon. Shading is SEM. **(B)** Responses of indicators in HEK293a cells to an AP burst waveform with spike widths of 1.0 ms at FWHM, and with 1 (top) to 4 spikes present (bottom). Waveforms are displayed in black on the far left. The voltage waveforms were previously recorded from CA1 mouse hippocampal pyramidal neurons at 37 ºC. V_m_ was –52 mV at the start of the displayed voltage segment. Each colored line is a single, unfiltered trace with no trial averaging or filtering applied.

**Figure S7.**
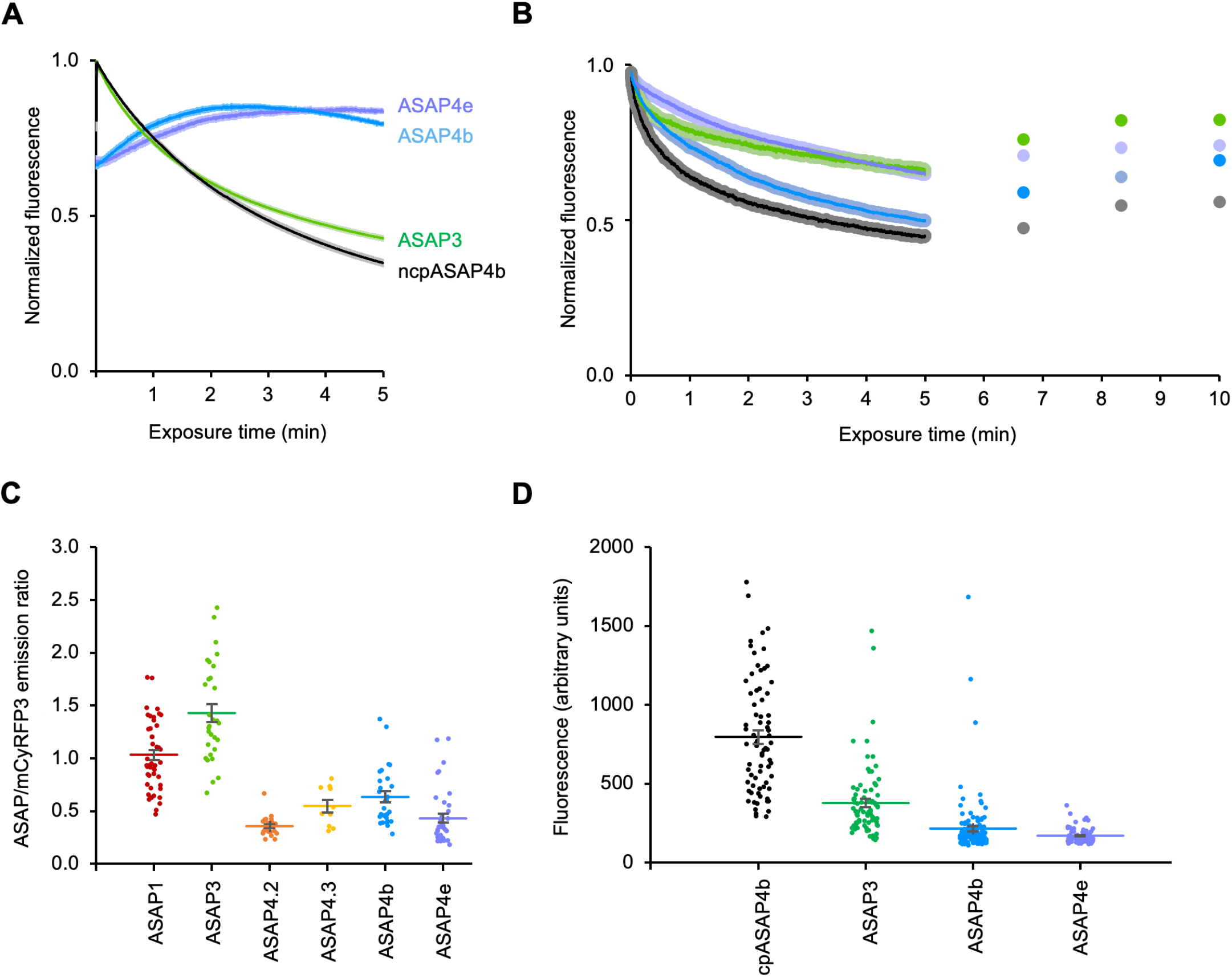
Brightness and Photobleaching of ASAP Indicators under 1-P and 2-P Imaging Modalities. **(A)** 1-P photobleaching curve over 5 minutes of continuous illumination in cultured HEK293-Kir2.1 cells, using blue light with a peak at 453 nm and an intensity of 50 mW/mm^2^. Notice the photoactivation present in ASAP4b and ASAP4e. ncpASAP4b n = 132 cells, ASAP3 n = 97 cells, ASAP4b n = 260 cells, ASAP4e n = 180 cells. Error bars are SEM. **(B)** 2-P photobleaching curves over 5 minutes in HEK293-Kir2.1 cells. The dots to the right of the curves represent where the curve average recovered to after 100, 200 and 300s, respectively. Cells were imaged under either 930nm light for ASAP4b and ASAP4e, or 940nm light for ASAP3 and ncpASAP4b, both at 960mW/mm^2^ which was the maximum power we could get from our setup. Recovery between 100 and 200 second pauses was significant across constructs (ASAP3 n = 84 cells, p 0.0189; ASAP4b n = 84 cells p = 0.0189; ASAP4e n = 76 cells p = 0.0419; ncpASAP4b n = 73 cells, p < 0.0001). However, none were significant between 200 and 300 seconds, with ASAP4b coming the closest (p = 0.0777), suggesting that after 200 seconds there are diminishing returns to keeping illumination off when allowing for recovery. Brown-Forsythe and Welch ANOVA used with Dunnett T3 test and p value corrected for multiple comparisons was used for all statistical tests. Shaded areas represent SEM. **(C)** ASAP variants and cytosolic mCyRFP2 were co-expressed in cultured hippocampal neurons at DIV 10 and imaged at DIV 12. The ratio of green to red brightness was plotted. ASAP3 was significantly brighter than all ASAP4 mutants, p = <0.0001 Dunnett’s multiple comparison test. ASAP4b was not significantly brighter than ASAP4e once multiple comparisons were accounted for, which brought the p-value to 0.0638 using Dunnett’s test. Error bars represent SEM. Sample sizes were as follows: ASAP1 (n = 48), ASAP3 (n = 30), ASAP4.2 (n = 23), ASAP4.3 (n = 10), ASAP4b (n = 28), and ASAP4e (n = 38). **(D)** 940-nm 2-P imaging of HEK293-Kir2.1 cells expressing either ncpASAP4b as a control (non-circularly permuted GFP fused to the ASAP4b VSD (n = 73), ASAP3 (n = 81), ASAP4b (n = 126) or ASAP4e (n = 76). Each data point is a cell, and the bar represents the mean value. Error bars represent SEM. All mutants were significantly different from each other with multiplicity adjusted p < 0.0001, except for ASAP4b vs ASAP4e which had multiplicity adjusted p = 0.8108 using Dunnett’s multiple comparison test.

**Figure S8.**
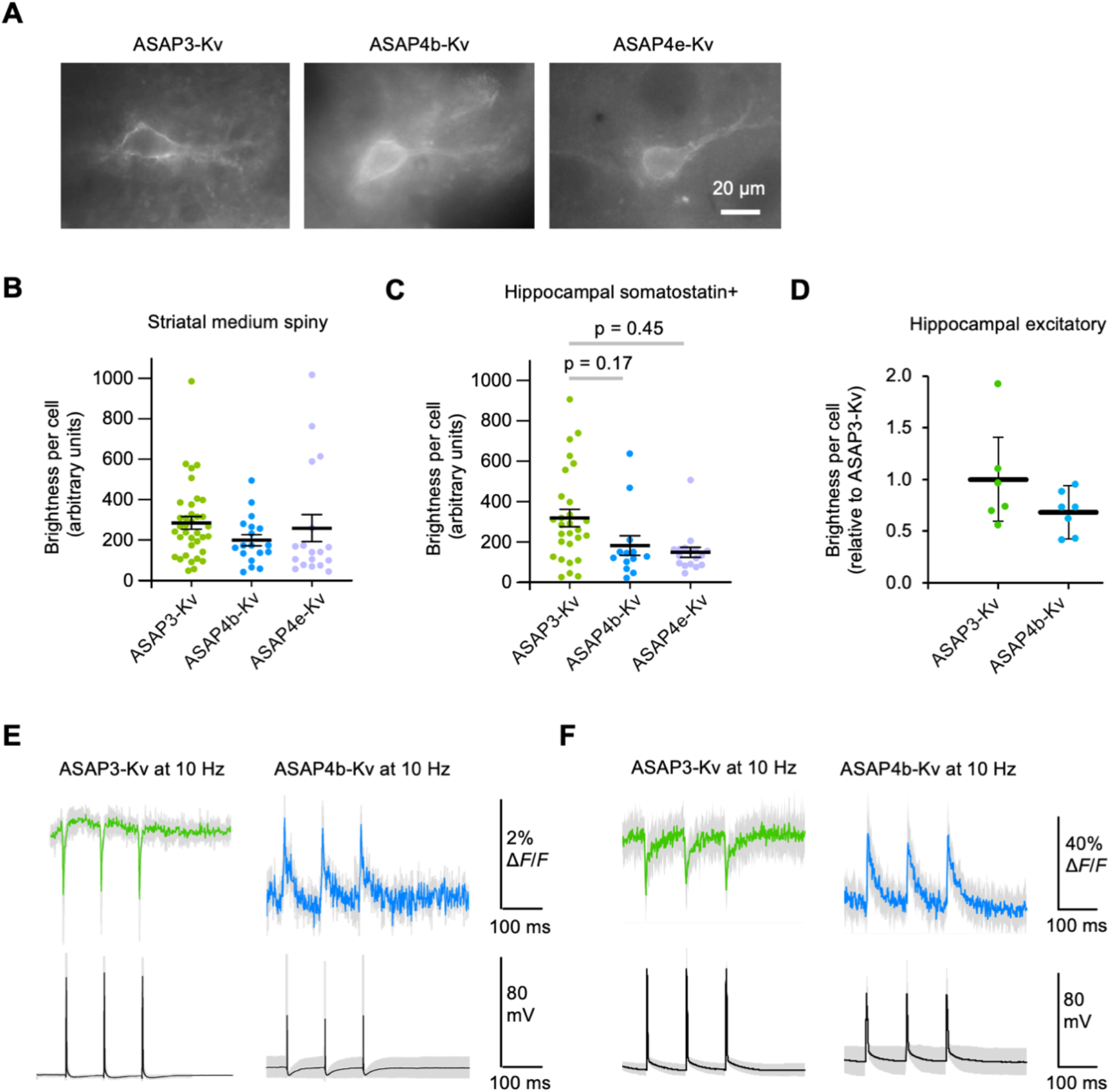
ASAP performance and brightness in brain tissue. **(A)** Examples of each indicator in acute striatal slice under widefield blue light illumination. Scale bar is 20μM. **(B)** Brightness values of striatal medium spiny neurons in acute coronal slice under blue light illumination. 6 cells were chosen at random from each mouse to match the mouse with the least number of cells, giving n = 36 cells for ASAP3-Kv, n = 18 cells for ASAP4b-Kv, and n = 18 cells for ASAP4e-Kv. Error bars are SEM. None of the differences in brightness were significant by Kruskal Wallis test. **(C)** The average brightness of the SOM+ hippocampal neurons recorded in vivo. ASAP3-Kv (green, n = 28), ASAP4b-Kv (blue, n = 13), and ASAP4e-Kv (purple, n = 17) are displayed. None of the differences were significant. **(D)** Acute slice 2-P (940nm) brightness. Background-subtracted fluorescence was measured across all Hippocampal CA1 pyramidal cells that were patch clamped, then normalized to the mean of the values from ASAP3-Kv mice. Bars represent the means of the brightness values across all mice for each indicator. Error bars show SEM, n = 6 cells for ASAP3-Kv and n = 7 cells for ASAP4b-Kv. Differences in brightness were insignificant by Mann Whitney test (p = 0.1678). **(E)** 1-P imaging of acute hippocampal slice cells expressing either ASAP3-Kv (n = 4 cells, green) or ASAP4b-Kv (n = 4 cells, blue) while being patch clamped in current clamp to evoke 10hz spike trains with current pulses, with 3 sweeps per cell. Below in black is the average voltage waveform recorded. Gray areas in all panels represents the SEM. **(F)** 2-P Imaging of acute hippocampal slice cells expressing either ASAP3-Kv (n = 6 cells, 21 sweeps total in green) or ASAP4b-Kv (n = 5 cells, 15 sweeps total in blue) while being patch clamped in current clamp to evoke 10hz spike trains with current pulses. Below in black is the average voltage waveform recorded. Gray shading is the SEM in all trace averages.

**Figure S9.**
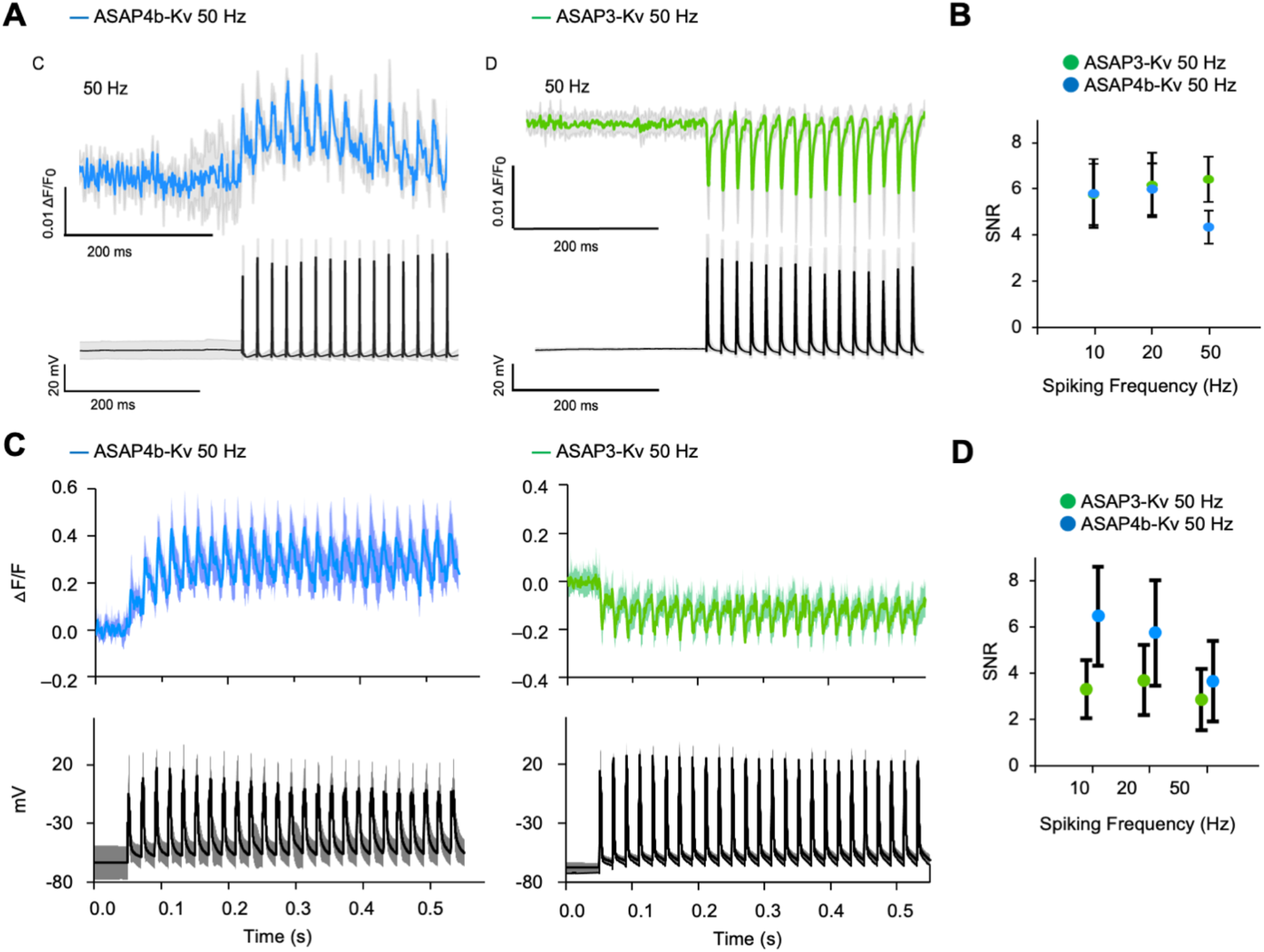
Detection of 50-Hz AP Trains in Acute Hippocampal Slice. **(A)** Top: Spikes were recorded under 1-P imaging at 1000hz The average unfiltered fluorescence trace of 50 Hz action potentials is shown, and was evoked during whole cell current clamp by brief current pulse injections in mouse hippocampal cells expressing ASAP4b-Kv or ASAP3-Kv. Bottom: The corresponding average voltage response from the same cells. The shaded regions represent the standard deviation, with n=4 cells for each. **(B)** The empirical signal-to-noise ratio for single spikes of ASAP3-Kv and ASAP4b-Kv at 10, 20 and 50 Hz as (ΔF/F0)/std(F0), from the same cells as (A). Error bars are Std. A Kruskal-Wallis test followed by Dunn’s multiple comparison test was used to compare ASAP4b-Kv to ASAP3-Kv within spiking frequency categories. Only 50hz was significantly different, with ASAP3-Kv having a higher SNR than ASAP4b-Kv (multiplicity adjusted p<0.0001). The same statistical test was done again, but this time comparing spiking frequencies within either ASAP3-Kv or ASAP4b-Kv. The ASAP3-Kv SNR at 10, 20 and 50hz were all not significantly different from eachother, while ASAP4b-Kv at 10 and 20hz were significantly higher than ASAP4b-Kv at 50hz with a multiplicity adjusted p-value of 0.0307 for 10hz vs 50hz, and 0.0002 for 20 vs 50 Hz. **(C)** Top: The average unfiltered fluorescence trace of 50hz evoked action potentials under a 2-P line scan imaging at 1000hz. Bottom: The corresponding average voltage response from the same cells. The shaded regions represent the standard deviation, with n = 3 cells with 15 trials total for ASAP4b-Kv, and n = 6 cells with 21 trials total for ASAP3-Kv. **(D)** Empirical SNR over the evoked firing rate for the 2-P imaging dataset in (C). Error bars are STD. A Kruskal-Wallis test followed by Dunn’s multiple comparison test was used to compare ASAP4b-Kv to ASAP3-Kv within spiking frequency categories, and to compare ASAP3-Kv and ASAP4b-Kv against themselves between spiking frequency categories. ASAP4b-Kv had a significantly higher SNR than ASAP3-Kv across all spiking frequency categories. Within ASAP3-Kv and ASAP4b-Kv, all comparisons were significant with multiplicity adjusted p < 0.0001, except for 10 vs 50 Hz for both indicators. For ASAP3-Kv, 10 vs 50 Hz was significantly different with multiplicity adjusted p = 0.0075.

**Figure S10.**
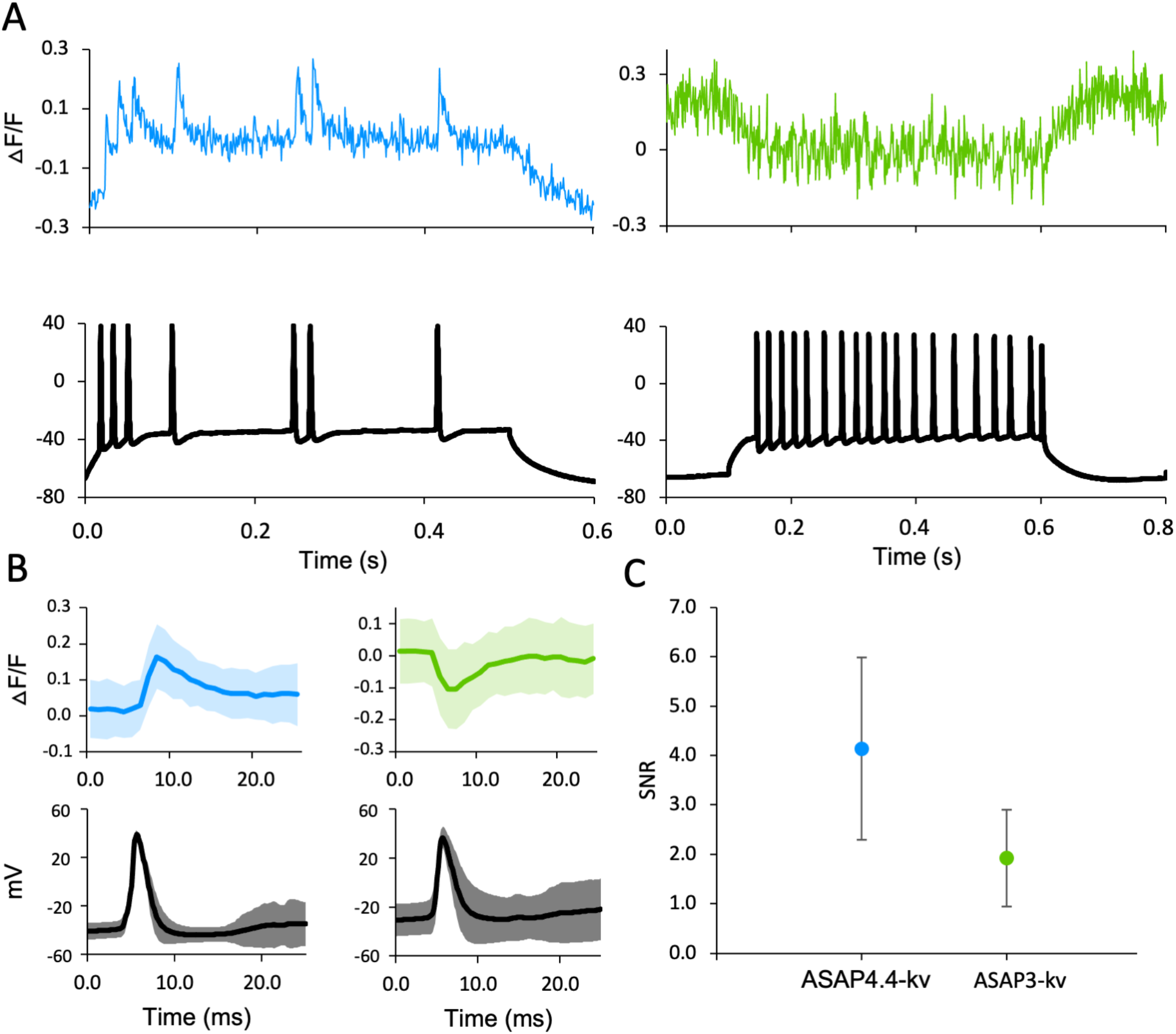
Current Step Evoked Spikes in Hippocampal CA1 Layer Pyramidal Neurons Under 2-P Imaging. **(A)** Top, examples of raw uncorrected fluorescence traces of ASAP4b-Kv (light blue) and ASAP3-Kv (green), recorded with 1000-Hz line scans of 2-P 940-nm excitation. Bottom, the corresponding electrophysiological traces for the raw fluorescence traces shown above. Spikes were evoked by applying a 100 pA current step, and then incrementing by 50–100 pA per step until the observed spikes began attenuating. **(B)** Shading for all sub-figures is STD. Top left figure: The average fluorescence response to all current step evoked spikes that attained at least 20mV in voltage for ASAP4b-Kv, n = 275 spikes. Top right figure: The same, but for ASAP3-Kv, n = 611 spikes. Bottom left figure: The average voltage trace for the ASAP4b-Kv fluorescence traces shown in the sub-figure above it. Bottom right figure: The average voltage trace for the ASAP3-Kv current evoked spike fluorescence traces shown above it. **(C)** Plot of the empirically determined SNR for the fluorescence traces in (B), defining noise as the std dev of the baseline brightness value before the start of the spiking protocol. Error bars are STD. ASAP4b-Kv (n = 275 spikes) had significantly higher SNR than ASAP3-Kv (n = 611 spikes) with p < 0.0001 using a Mann-Whitney test.

**Figure S11.**
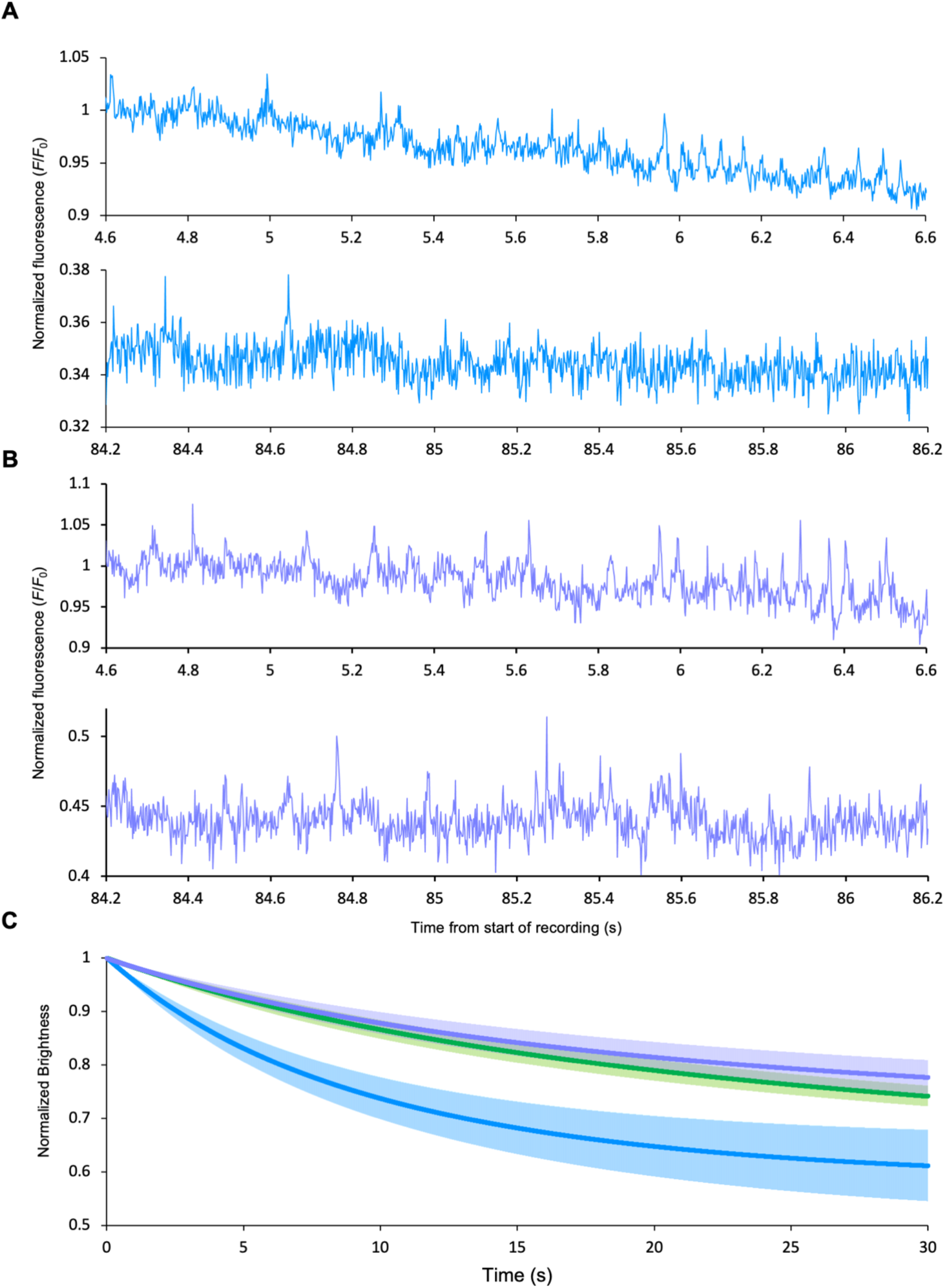
In Vivo Photobleaching Under 1-P Illumination in Hippocampal Sst+ Interneurons. **(A)** An example trace of ASAP4b-Kv in vivo in an Sst+ interneuron of the Hippocampus. The upper panel shows the first 2 seconds of recording after the first 4.6 seconds of illumination, while the bottom panel shows the recording up to 86.2 seconds later, but normalized to the brightness in the initial recording, so the photobleaching can be seen at 247mW/mm^2^. The cell is displayed at 500hz. The trace has not been corrected for power, or filtered in any way. Cells were imaged for 11 seconds with continuous illumination during a running bout, then 8.1 seconds of darkness when the running bout ended. The dark periods were not included in the time axis here, as the illumination periods were concatenated. **(B)** An example trace of ASAP4e-Kv in vivo in an Sst+ interneuron. The panels show examples in the same style as (A), except for the power, which was 95mW/mm2, and is shown at 500 hz. The trace has not been corrected for power, or filtered in any way. **(C)** In vivo 1-photon photostability. n = 6 cells for ASAP3-Kv (green), n = 7 cells for ASAP4b-Kv (blue), n = 8 cells for ASAP4e-Kv (purple). Error bars are the SEM. All traces were normalized to 100mW to correct for differences in power, under linear assumptions e.g. 2-fold power increase causes 2-fold faster photobleaching along a monoexponential curve, which was implemented by changing the slope of the log of the data.

## METHODS

### Plasmid Construction

For transfection into HEK293-Kir2.1 Cells for electrical screening, the parent ASAP2f L146G S147T R414Q was subcloned into pcDNA3.1 with a CMV enhancer and promoter and bGH poly(A) signal. All plasmids were made by standard molecular biology techniques with all cloned fragments confirmed by sequencing (Sequetech). PCR reactions were carried out to generate PCR product libraries using standard PCR techniques, which were used to directly transfect HEK293-Kir2.1 cells with the linear product using lipofectamine 3000 (Thermo Fisher Scientific). For patch clamp characterization in HEK293A cells, all voltage indicators were subcloned into a pcDNA3.1/Puro-CAG vector between NheI and HindIII sites (*20*).

For in vitro characterization in cultured neurons, acute hippocampal slice and in vivo hippocampal imaging, ASAP variants were subcloned into pAAV.hSyn.WPRE. These were then midi-prepped and packaged into adeno-associated virus 8 (AAV8) by the Neuroscience Gene Vector and Virus Core of Stanford University. For somatic targeting in acute slice and in vivo, the C-terminus of ASAP4 variants was attached to the C-terminal cytoplasmic segment of the Kv2.1 potassium channel (*22*), that we previously used to restrict ASAP3 to the soma, axon and proximal dendrites (*39*). For EF1α-driven ASAP expression, viral constructs were generated by modifying published methods (*42*) using GateWay recombination and Gibson assembly, and produced in-house as previously described (*39, 49*). These were then packaged into AAV8 capsids by the Neuroscience Gene Vector and Virus Core at Stanford University.

### Cultured Cell Lines

All cell lines were maintained in a humidified incubator at 37°C with 5% CO_2_. For electrical screening the previously described HEK293-Kir2.1 cell line (*47*) was maintained in high-glucose DMEM, 5% FBS, 2 mM glutamine, and 500 μg/mL geneticin (Life Technologies). For patch-clamp recordings to measure ASAP responsivity and kinetics, HEK293A cells were cultured in high-glucose Dulbecco’s Modified Eagle Medium (DMEM, Life Technologies) with 5% fetal bovine serum (FBS, Life Technologies) and 2 mM glutamine (Sigma-Aldrich).

### Cultured Neurons

Hippocampal neurons were isolated from embryonic day 18 Sprague Dawley rat embryos of both sexes. Procedures were carried out in compliance with the rules of the Stanford University Administrative Panel on Laboratory Animal Care.

### Viruses

AAV9-CamKII-Cre was obtained from the Penn Vector Core. AAV1-Syn-NES-jRGECO1a, ≥1×10^11^ vg/mL, Addgene #100854, and AAV1-syn-NES-jRGECO1b-WPRE-SV40, AddGene #100857, were ordered from Addgene. All ASAP viruses were produced by the Stanford Neuroscience Gene Vector and Virus Core facility, and by the lab of Sui Wang at Stanford University.

For the acute slice brightness comparisons, viruses were diluted in phosphate-buffered saline (PBS) until the following titers were reached: AAV9-CamKII-cre at a titer of 5.6×10^9^ vg/mL, and all three ASAP viruses at a titer of 3.45×10^11^ vg/mL. For the 2-P imaging while patch clamping evoked spikes in hippocampal slice work, AAV8-syn-ASAP3/4b-Kv was injected at a titer between 1.21×10^12^ and 5×10^12^ vg/mL, and AAV1-syn-jrGECO1b was injected at a titer between 1.3×10^12^ and 5×10^12^ vg/mL, but we were not able to image the jRGECO1b well, so it was excluded from the analysis. For the 1-P imaging of patch clamping in hippocampal slice and evoking spikes, AAV8-syn-ASAP4b-Kv and AAV8-syn-ASAP3-Kv were injected at titers approximately between 1×10^12^ and 1×10^13^ vg/mL. For the in vivo 2-P imaging of ASAP4e and jRGECO1a in V1, AAV8-Syn-ASAP4e-Kv at a titer of 4.66×10^11^ vg/mL was co-injected with AAV1-Syn-NES-jRGECO1a at a titer of approximately 1×10^11^ vg/mL. For the in vivo 1-P imaging of hippocampus during running, either AAV8-ef1α -DiO-ASAP3-Kv at a titer of 2.35×10^12^ vg/mL, AAV8-ef1α -DiO-ASAP4b-Kv at a titer of 2.36×10^12^ vg/mL, or AAV8-ef1α -DiO-ASAP4e-Kv at a titer of 3.45×10^11^ vg/mL was injected. For the in vivo 2-P imaging of the hippocampus during a spatial navigation task, AAV8-syn-ASAP4b-Kv at a titer of 1.17 × 10^12^ vg/mL was co-injected with AAV1-syn-NES-jRGECO1b-WPRE-SV40 at a titer of 1.3×10^13^.

### Animals

For the 1-P acute slice brightness comparisons, 2-P acute slice patch clamping experiments and in 2-P in vivo imaging experiments, adult male and female wild-type C57BL/6 were used. All mice were housed in standard conditions (up to five animals per cage, 12-hour light/dark cycles with the light on at 7 a.m., with water and food ad libitum). Day-18 Sprague Dawley rat embryos were used for hippocampal tissue for neuronal culture. All protocols were approved by the Stanford Institutional Animal Use and Care Committee.

For the in vivo fly imaging, ASAP4b was cloned into the pJFRC7-20XUAS vector (*28*) using standard molecular cloning methods (GenScript Biotech), and then inserted into the attP40 phiC31 landing site by injection of fertilized embryos (BestGene). We used the cell type-specific driver 21D-Gal4 (*33*) to express ASAP2f and ASAP4b in L2 cells. We imaged the axon terminals of L2 cells in its medulla layer M2 arbors. The genotypes of the imaged flies in Fig. S4 were: L2>>ASAP2f: +; UAS-ASAP2f/+; 21D-Gal4/+. L2>>ASAP4b: yw/+; UAS-ASAP4b/+; 21D-Gal4/+. All protocols were approved by the Stanford Institutional Animal Use and Care Committee.

In vivo 1-P imaging experiments were conducted at the University of California at Los Angeles (UCLA), where adult (11-23 week old) Sst-IRES-cre knock-in adult male and female mice were used for all experiments. All animals were group housed (2-5 per cage) on a 12 h light/dark cycle. All experimental protocols were approved by the Chancellor’s Animal Research Committee of the University of California, Los Angeles, in accordance with NIH guidelines.

Animals used in acute slice 1-P patch clamping experiments were male and female wild-type mice aged 20-40 days. All mice used were maintained on a 12 hr light/dark cycle, with food and water available ad libitum and in accordance with the Institutional Animal Care and Use Committee of Columbia University.

For the co-imaging of ASAP4e and jRGECO1a in V1, all animal experiments were conducted according to the National Institutes of Health guidelines for animal research. Procedures and protocols on mice were approved by the Animal Care and Use Committee at the University of California, Berkeley.

## Method details

### Cell screening

Cell screening was the same as previously described in Villette et. al. 2019. Briefly, HEK293-Kir2.1 cells were plated in 384-well plates (Grace Bio-Labs) on conductive glass slides (Sigma-Aldrich). Cells were transfected with PCR-generated libraries in 384-well plates with Lipofectamine 3000 (∼100 ng DNA, 0.4 μL p3000 reagent, 0.4 μL Lipofectamine) followed by a media change 4–5 hours later, with imaging done 2 days post-transfection in Hank’s Balanced Salt solution (HBSS) buffered with 10 mM HEPES (Life Technologies). Cells were imaged at room temperature on an IX81 inverted microscope fitted with a 20× 0.75-numerical aperture (NA) objective (Olympus). A 120-W Mercury vapor short arc lamp (X-Cite 120PC, Exfo) served as the excitation light source. The filter cube set consisted of a 480/40-nm excitation filter and a 503-nm long pass emission filter. ASAP libraries were screened at room temperature with the operator locating the best field of view and focusing on the cells. A single field of view was imaged for a total of 5 s, with a 10-μs 150-V square pulse applied near the 3-s mark. Fluorescence was recorded at 100 Hz (10-ms exposure per frame) by an ORCA Flash4.0 V2 C11440–22CA CMOS camera (Hamamatsu) with pixel binning set to 4 × 4. Mutants were screened at least three times.

### Whole cell patch clamping and imaging of hek293a cells

Methods were the same as those described in Villette et. al., 2019, and modifications are described as follows. Briefly, ASAPs and upward Ace2N-mNeon variants were subcloned into a pcDNA3.1/Puro-CAG vector, and cells were transfected using Lipofectamine 3000 (400 ng DNA, 0.8 μL P3000 reagent, 0.8 μL Lipofectamine) per manufacturer’s recommended instructions. After plating the cells on 12 mm glass coverslips (Carolina Biological), the cells were patch clamped 24 hours after transfection. Patch-clamp experiments were mainly done as previously described in Villette et. al., 2019. Signals were recorded in voltage-clamp mode with a Multiclamp 700B amplifier and using pClamp software (Molecular Devices). Fluorophores were illuminated at ∼4.3 mW/mm^2^ power density at the sample plane with an UHP-Mic-LED-460 blue LED (Prizmatix) passed through a 484/15-nm excitation filter and focused on the sample through a 40 × 1.3-NA oil-immersion objective (Zeiss). Emitted fluorescence passed through a 525/50-nm emission filter and was captured by an iXon 860 electron-multiplied charge-coupled device camera (Oxford Instruments) cooled to −80 °C. For all experiments fluorescence traces were corrected for photobleaching by dividing the recorded signal by a mono-exponential fit to the data. To characterize steady-state fluorescence responses, cells were voltage-clamped at a holding potential of –70 mV, then voltage steps of 1 s duration between –120 and +120 mV were imposed. The specific steps used were -120mV, -100mV, -80mV, -60mV, -40mV, -20mV, 0mV, 30mV, 50mV, 70mV, 90mV, and 120mV. To obtain complete voltage-tuning curves, steps between –200 and +120 mV were imposed. The specific steps were -200mV, -180mV, -160mV, -140mV, -120mV, -100mV, -80mV, -60mV, -40mV, -20mV, 0mV, 30mV, 50mV, 70mV, 90mV, and 120mV. When voltage-clamp could not be imposed on some cells at the highest potentials, that step was excluded from data analysis. A scaled action potential waveform (FWHM=3.5ms, -70mV to +30mV) recorded from a cultured hippocampal neuron was included before each step to estimate the fluorescence change of the indicators to action potential waveform. To characterize kinetics of ASAP indicators at both room temperature and 37°C, images were acquired at 2.5 kHz from an area cropped down to 64 × 64 pixels and further binned to comprise only 8 × 8 pixels. Fluorescence was quantified by averaging pixels with signal change in the corresponding cell shape after subtracting by the background pixels. Command voltage steps were applied for 1 s, and a double exponential or single exponential fit was applied to a 60-ms interval from the starting point of the onset and offset of voltage steps using MATLAB software (MathWorks). To characterize ASAP and upward Ace2N-mNeon variants on simulated AP burst waveforms (2ms and 4ms at FWHM) at room temperature, images were acquired at 1000Hz from a 64 × 64 pixel FOV, then binned using 4 × 4 pixel binning. To characterize ASAP variants on square pulses (1ms, 2ms and 4ms, -70mV to +30mV) and AP burst waveforms (1.0 ms at FWHM) at 37 °C, images were acquired at 2500Hz from a 64 × 64 pixel FOV, then binned using 8 × 8 binning. For more details, please see Villette et. al., 2019.

### Brightness comparison of ASAP indicators in vitro

Cultured hippocampal neurons were transfected at 10 DIV via calcium phosphate with pcDNA3.1-ASAPx-Kv-mCyRFP2 variants. Cells were imaged at 12 DIV on an inverted confocal microscope (Olympus U-TB190) with a 20X objective (Olympus, NA 0.75). Images were taken using an Andor EMCCD897 camera with Andor IQ2.0 software. ASAPs and mCyRFP2 were excited with a 488 nm laser and fluorescence were collected via 514/30 nm and 607/36 nm filters, respectively. To correct crosstalk between the two channels, we constructed a colorless version of ASAP4b-Kv-mCyRFP2 with a Y312A mutation, which corresponds to the Y66A mutation in GFP. The fluorescence signal collected in the green channel was 3.28% of that of the red channel (n = 58 neurons) and was then subtracted for all the other variants. For a control comparison, we also changed the cpsfGFP part of ASAP4b into a non-circular-permuted version. The artificial cerebral spinal fluid (ACSF) recipe used was as follows: NaCl (150 mM), KCl (3 mM), CaCl_2_ (3 mM), MgCl_2_ (2 mM), HEPES (10 mM), and Glucose (5 mM) in deionized (DI) H_2_O.

### Primary neuronal culture and transfection

Methods used here were the same as in Villette et. al., 2019. Briefly, hippocampal neurons were isolated from embryonic day 18 Sprague Dawley rat embryos by dissociating them in RPMI medium containing 5 units/mL papain (Worthington Biochemical) and 0.005% DNase I at 37º C and 5% CO_2_ in air. Neurons were plated on washed 12-mm No.1 glass coverslips pre-coated overnight with > 300-kDa poly-D-lysine hydrobromide (Sigma-Aldrich). Cells were plated for several hours in Neurobasal media with 10% FBS, 2 mM GlutaMAX, and B27 supplement (Life Technologies), then the media was replaced with Neurobasal with 1% FBS, 2 mM GlutaMAX, and B27. Half of the media was replaced every 3–4 days with fresh media without FBS. 5-Fluoro-2’-deoxyuridine (Sigma-Aldrich) was typically added at a final concentration of 16 μM at 7–9 DIV to limit glial growth. Hippocampal neurons were transfected at 9–11 DIV using Lipofectamine 2000 (Life Technologies, 100–300 ng of indicator DNA with empty pNCS vector together as 500 ng DNA in total, 1 μL lipofectamine 2000, 200 μL Neurobasal with 2mM GlutaMax). See Villette et. al. 2019 for more details.

### High resolution cultured hippocampal neuron imaging

Cultured hippocampal neurons were transfected at 9–11 DIV via lipofectamine 3000 (Invitrogen) with pAAV-hSyn-ASAPx-WPRE and pAAV-hSyn-Ace2-D92N-E199V-mNeon-ST-WPRE (upward Ace2N-mNeon) variants. Cells were imaged at 12–20 DIV on an inverted confocal microscope (Zeiss Axiovert 200M) with a 40X objective (Zeiss, NA 1.20). ASAPs and upward Ace2N-mNeon were excited with 488/30 nm filters and fluorescence were collected via 531/40-nm filters by a 120-W Mercury vapor short arc lamp (X-Cite 120PC, Exfo). Images were taken using an ORCA Flash4.0 V2 C11440-22CA CMOS camera (Hamamatsu) with Micro-manager software. Exposure time was 1 second per frame, with frames collected as 2048×2048 16-bit images. (*15*)

### 1-P photobleaching in hek293-kir2.1 cells

Illumination was provided with a blue LED light source (peak at 453 nm, UHP-F3-455 with LLG-3 light guide, Prizmatix). HEK293-Kir2.1 cells plated on poly-D-lysine coated 12-mm diameter coverslips (0.13–0.17 mm thickness; Carolina Biological) were transiently transfected using Lipofectamine 3000 (Life Technologies) at 60– 80% cell confluency by using plasmids with a CMV promoter (200–300 ng DNA, 0.8 μL P3000, and 0.8 μL Lipofectamine 3000). The transfected cells were then imaged 1–3 days post-transfection. During imaging, cells were bathed in Hank’s Balanced Salt Solution (HBSS) with 10 mM HEPES, which was pumped with a Masterflex HV-77122-24 Variable-Speed pump. Light was filtered with a U-N41017 EN GFP filter cube (Chroma) through a 40x water immersion objective (Olympus LUMPlanFL 40x), before being imaged with an Orca Flash 4.0LT camera being controlled by HCImage Live (Hammamatsu).

### 2-P photobleaching in HEK293-Kir2.1 cells

Cultured cells were the same as those listed under *1-P photobleaching in HEK293-Kir2*.*1 cells* above. A custom-built 2-P microscope system using a tunable 2-P laser (InSight X3, Spectra-Physics, Santa Clara, CA) was modulated with a Pockel’s cell (350-80, Conoptics, Danbury, CT). The laser directed with a 12.0-kHz resonant scanner (*19*) and was focused through a 40×, 0.8-NA objective (Nikon, Japan). Data were collected through a PMT (H10770PA-40, Hamamatsu, Japan), after being filtered for green light (ET525/50M, Chroma Technology).

### 2-P in vivo imaging of *Drosophila*

Flies were mounted, dissected to expose the brain, perfused with a saline-sugar solution, and imaged for up to 1h, as previously described (*8, 46*). Neurons were excited at 920 nm with 5–15 mW of total power, and photons were collected with a 525/50-nm filter. Data was collected at an 82.4 Hz frame rate with 200×20-pixel frames using a 15X digital zoom, using bidirectional scanning. We used a Leica TCS SP5 II two-photon microscope with a Leica HCX APO 20×/1.0-NA water immersion objective (Leica) and a pre-compensated Chameleon Vision II femtosecond laser (Coherent, Inc.).

### Visual stimulation

Visual stimuli were generated with custom-written software using MATLAB (MathWorks) and presented using the blue LED of a DLP Lightcrafter 4500 (Texas Instruments) projector in Pattern Sequence mode. The stimulus was refreshed at 300 Hz and utilized 6 bits/pixel, allowing for 64 distinct luminance values. The stimulus was projected onto a 9 × 9-cm rear-projection screen positioned approximately 8 cm anterior to the fly that spanned approximately 70° of the fly’s visual field horizontally and 40° vertically. A small square was also simultaneously projected onto a photodiode (Thorlabs, SM05PD1A) configured in a reversed-biased circuit. The stimulus was filtered with a 482/18-nm bandpass filter so that it could not be detected by the microscope PMTs. The radiance at 482 nm was approximately 78 mW sr^−1^ m^−2^.

The imaging and the visual stimulus presentation were synchronized using triggering functions provided by the LAS AF Live Data Mode software (Leica) as well as the signal from the photodiode directly capturing projector output. A Data Acquisition Device (NI DAQ USB-6211, National Instruments) connected to the computer used for stimulus generation was used to acquire the photodiode signal, generate a trigger signal at the beginning of stimulus presentation, and acquire the trigger produced by the LAS software at the start of each imaging frame. This allowed the imaging and the stimulus presentation to initialize in a coordinated manner and ensured that stimulus presentation details were saved together with imaging frame timings (in MATLAB .mat files) to be used in subsequent processing. Data was acquired on the DAQ at 5000 Hz.

The visual stimuli used were 300-ms search stimuli: alternating full contrast light and dark flashes, each 300 ms in duration, were presented at the center of the otherwise dark screen. The stimulus was such that from the perspective of the fly, the flashing region was 8° from each edge of the screen. In subsequent analysis, the responses to this stimulus were used to select ROIs with receptive fields located at the center of the screen instead of at the edges. This stimulus was presented for 5000 imaging frames (61 s) per field of view. Single 20-ms light and dark flashes, with 500-ms of gray between the flashes, were presented over the entire screen. The light and dark flashes were randomly chosen at each presentation. The Weber contrast of the flashes relative to the gray was 1. This stimulus was presented for 10,000 imaging frames (122 s) per field of view.

### In vivo imaging data analysis

The acquired time series were saved as .lif files and read into MATLAB using Bio-Formats (Open Microscopy Environment). Each time series was aligned in x and y coordinates by maximizing the cross-correlation in Fourier space of each image with a reference image (the average of the first 30 images in the time series). For each time series, ROIs around individual arbors were selected by thresholding the series-averaged image with a value that generates appropriate ROIs, and then splitting any thresholded ROIs consisting of merged cells and/or adding ROIs that were missed by the thresholding.

The ΔF/F for each time series was calculated after subtracting the background intensity and correcting for bleaching as previously described (*8, 46*). Time series with uncorrected movement, which was apparent as irregular spikes or steps in the ΔF/F traces that were coordinated across ROIs, were discarded. The stimulus-locked average response was computed for each ROI by reassigning the timing of each imaging frame to be relative to the stimulus transitions (gray to light or gray to dark) and then computing a simple moving average. The averaging window was 8.33 ms and the shift was 8.33 ms, which effectively resampled our data from 82.4 Hz to 120 Hz.

As the screen on which the stimulus was presented did not span the fly’s entire visual field, only a subset of imaged ROIs experienced the stimulus across approximately the entire extent of their spatial receptive fields. These ROIs were identified based on having a response of the appropriate sign to the 300-ms search stimulus. ROIs lacking a response to these stimuli or having one of the opposite sign were not considered further. The peak response to each flash (peak Δ*F*/*F*) was the Δ*F*/*F* value farthest from zero in the expected direction of the initial response (depolarization or hyperpolarization). The time to peak (t_peak_) was the time at which this peak response occurred, relative to the start of the light or dark flash. Pairwise Student’s t-tests were performed.

### Acute slice brightness comparisons

Viruses were diluted with PBS until the following titers were reached: 3.45×10^11^ for AAV8-EF1α-FLex-ASAP3/4.4/4.5 and 5.6×10^9^ AAV9-CamkII-cre, which was obtained from the UPenn viral core, and is plasmid #105558 on Addgene. Mice were injected between p40 and p56. Each mouse was injected with 950-1000 nL at a flow rate of 200nL/min. Acute slices were obtained 28-36 days post injection. Injections were performed at the following coordinates relative to Bregma: AP: +1.2mm; ML: –2mm; DV: –3.2 to –2mm, placing them into the dorsal striatum. Light was delivered to and received from the samples via a U-N41017 EN GFP filter cube (Chroma), through a 40x water immersion objective (Olympus LUMPlanFL 40x). To control for fluctuations in illumination intensity, videos of 50 frames were recorded at 33hz with a 30ms exposure time, and the average taken. Imaging was done with an orca-Flash4.0 LT. ROIs for the cells were drawn in ImageJ (National Institute of Health), taking the mean pixel value within the ROI. Background was determined as the tissue near the cell without any cell debris present, and the mean was taken and subtracted out to give the final brightness value for a cell.

### 1-P imaging while patch clamping evoked spikes in hippocampal slice

Mice aged 20–40 days received injections with AAV virus carrying ASAP3-Kv or ASAP4b-Kv. 4-7 days post injection, animals were anesthetized with 5% isoflurane and acute coronal slices were prepared as previously described (*25*). After achieving whole-cell recording configuration, action potentials were evoked in current clamp mode using 1 ms injections of 1 pA. Images were simultaneously recorded at ∼1000 frames per second using a coolLED-pe300 light source (coolLED LTD, Andover UK) and a photometrics prime-95B camera (Teledyne Photometrics, Tuscon AZ) with an Olympus LUMPLFLN 40×W, 0.8 NA.

### 2-P imaging while patch clamping evoked spikes in hippocampal slice

Mice between p28 and p60 were injected with a mixture of AAV8-syn-ASAP3/b-Kv.WPRE and AAV1-syn-jRGECO1b-WPRE totalling 1 uL in volume. Injections were performed unilaterally into the right hemisphere under either ketamine/xylazine or isofluorane anesthesia. AAV8-syn-ASAP3/4b-Kv was injected at a titer between 1.21×10^12^ and 5×10^12^. AAV1-syn-jRGECO1b was injected at a titer between 1.3×10^12^ and 5×10^12^. The mixture was injected at 100nL/min at the following coordinates from bregma: AP: -1.5mm. ML: -1.5mm. DV: -1.5 to -1.3mm through a glass micropipette (VWR) pulled with a long narrow tip (size ∼10–20 μm) using a micropipette puller (Sutter Instrument). The pipette was gently withdrawn 5 min after the end of infusion and the scalp was sutured. Mice were sliced 4–8 weeks after AAV injection. Coronal brain slices (300 μm) containing the dorsal striatum were obtained using standard techniques (*12*). Briefly, animals were anesthetized with isoflurane and decapitated. The brain was exposed and chilled with ice-cold artificial cerebrospinal fluid (ACSF) containing 125 mM NaCl, 2.5 mM KCl, 2 mM CaCl2, 1.25 mM NaH2PO4, 1 mM MgCl2, 25 mM NaHCO3, and 15 mM D-glucose (300–305 mOsm). Brain slices were prepared with a vibrating microtome (Leica VT1200 S, Germany) and left to recover in ACSF at 34°C for 30 min followed by room temperature (20–22 C) incubation for at least additional 30 min before transfer to a recording chamber. The slices were recorded within 5 hours after recovery. All solutions were saturated with 95% O2 and 5% CO2 (Carbogen) in ACSF, which was pumped out of the recording chamber using a Masterflex HV-77122-24 pump. Hippocampal CA1 layer pyramidal neurons were visualized with infrared differential interference contrast (DIC) illumination and ASAP3/4b-Kv-expressing neurons were identified with epifluorescence illumination on a BX-51 microscope equipped with a 60× 1.0-NA water-immersion objective and DIC optics (Olympus) and a Lambda XL arc lamp (Sutter Instrument). Whole-cell current-clamp recording was performed with borosilicate glass microelectrodes (3–5 MΩ) filled with a K^+^ based internal solution (135 mM KCH3SO3, 8.1 mM KCl, 10 mM HEPES, 8 mM Na2-phosphocreatine, 0.3 mM Na2GTP, 4 mM MgATP, 0.1 mM CaCl2, 1 mM EGTA, pH 7.2–7.3, 285–290 mOsm). Access resistance was compensated by applying bridge balance. To induce firing, 1 ms pulses of 2-nA current were injected to induce spiking at different frequencies (10, 20 or 50, Hz). Recordings were obtained with a Multiclamp 700B amplifier (Molecular Devices) using the WinWCP software (University of Strathclyde, UK). Signals were filtered with a Bessel filter at 2 kHz to eliminate high frequency noise, digitized at 10 kHz (NI PCIe-6259, National Instruments). Two-photon imaging was performed with a custom built two-photon laser-scanning microscope as described previously (*7*), equipped with a mode-locked tunable (690–1040 nm) Mai Tai eHP DS Ti:sapphire laser (Spectra-Physics) tuned to 940 nm. The jRGECO signal in the red channel was ultimately not included because this setup was not able to obtain jRGECO signals at 940nm, and did not have enough power at higher wavelengths to obtain a good signal in the red channel.

ASAP signals were acquired by a 1-kHz line scan across the membrane region of a cell. Signals recorded along each line were integrated to produce a fluorescence trace over time, and a region of the line scan corresponding to the background beyond the cell membrane was chosen as the background value, and subtracted from the integrated membrane region. All traces were then normalized to 1 by either dividing by a monoexponential, or dividing by the first 0.5 s of data where no activity was present, if there was no photobleaching present and the exponential fit was not working well.

For the current steps, cells were patch clamped in current clamp mode, and taken through steps starting at 100pA, and increasing by 50 to 100pA until spiking was elicited, and then continued until spikes began to attenuate due to the depolarization blockade. Spikes were included in the SNR comparisons and average waveforms if the corresponding electrophysiological waveform crossed 20 mV in height at its peak. This excluded attenuated spikes from the analysis. Once detected in the data, spikes were aligned by taking the derivative of the waveform, and aligning the individual spikes by the peak of the derivative during the onset (where the onset of the voltage waveform is the steepest).

For the SNR calculation, the formula (Δ*F*/*F*_0_)/STD(*F*_0_) was used. *F*_0_ was determined by using the value of the monoexponential or linear fit right before the spike occurred in the raw data. For taking the standard deviation, the 50 points immediately preceding the detected spike peak in the raw data were used. Δ*F* is the maximum value attained by each spike after the trace was normalized to 1 by dividing by a monoexponential fit to the trace. The data points ± 2 ms of the peak of the average fluorescence waveform were searched to find the peak of single spikes, in order to account for timing jitter in the recordings.

### In vivo 2-P imaging of ASAP4e and jRGECO1a in the visual cortex

A wild-type (Jackson Laboratories, Black 6, stock no. 000664) mouse was used for simultaneous calcium and voltage imaging. The cranial window implantation procedure used has been described previously (*36*). In brief, the 3-month-old mouse was anesthetized with 1–2% isoflurane in O2 combined with the analgesic buprenorphine and head-fixed in a stereotax. A craniotomy was made over left V1, followed by pipet injection of 200nL of viral solution (3:1 ratio of AAV8-Syn-ASAP4e-Kv, 6.21×10^11^ vg/mL and AAV1-Syn-NES-jRGECO1a, ≥ 1×10^11^ vg/mL, Addgene #100854) at 300 μm below the exposed brain surface. A glass window was embedded and sealed in the craniotomy. A stainless-steel head-bar was firmly attached to the skull with dental acrylic. The implanted mouse was provided with the post-operative analgesic Meloxicam for 2 days and allowed to recover for 2 weeks prior to imaging experiments.

Imaging was performed while the mouse was head-fixed and awake. A modified, commercially available two-photon microscope was used (*13*). A titanium-sapphire laser (Chameleon Ultra II, Coherent Inc.) was used as the two-photon excitation source, and a wavelength of 1000 nm was chosen to excite both ASAP4e-Kv and jRGECO1a fluorescent proteins. Post-objective powers of 18–31mW were used for imaging of depths 140–185 μm below brain surface. Fields of view from 32×32 μm to 240×240 μm were imaged at a pixel size of 0.25×0.25 μm/pixel, resulting in frame rates varying from 15-99Hz. Images from two fluorescence emission channels (green: ASAP4e-Kv, red: jRGECO1a) were collected simultaneously. Image time stacks were first registered with rigid motion correction (NoRMCorre, MATLAB), then single neuron time traces were extracted by averaging the signal within hand-drawn ROIs using Fiji (ImageJ).

### In vivo 1-P imaging of hippocampus during running

Adult (11–23 weeks old) Sst-IRES-Cre mice (2 male, 5 female) were injected with 500 nL of either AAV8-EF1α-DiO-ASAP3-Kv (2 mice) at a titer of 2.35×10^12^, AAV8-ef1α-DiO-ASAP4b-Kv (2 mice) at a titer of 2.36×10^12^, or AAV8-EF1α-DiO-ASAP4e-Kv (4 mice) at a titer of 3.45×10^11^, in the right dorsal CA1 area at 60 nL/min. 455-nm light was used as it has been shown to increase photostability over 470 nm in ASAP family GEVIs (*44*). We kept maximum LED power at 5 mW total and 25 0mW/mm^2^ or less, as we found, and it has been previously shown, that sustained power levels above this resulted in tissue damage (*32*). Data was acquired at 1000 Hz while the mice ran on a wheel, in 11-sbehavioral intervals. In between each 11 second recording bout was an 8.1-s pause. All surgical procedures, injection site, postoperative and training protocols are the same as those described in (*37*).

Animals were imaged using a custom-built, high speed single-photon epi-fluorescent microscope. Photoexcitation was provided via a fiber-coupled LED (Thorlabs, M455F3) with a center wavelength of 455nm (*44*). Excitation light is collimated after a 2-m long, 400-μm core multi-mode fiber optic patch cord (Thorlabs, M28L02) and expanded using a Keplerian telescope. The expanded beam is passed through a spectral excitation filter (Thorlabs, MF455-45), and reflected off of a long-pass dichroic mirror (Thorlabs, MD480) before teaching a 16×/0.8NA water-immersion objective (Nikon, CFI75 LWD 16X W). The expander is used to generate a localized excitatory spot ∼165 μm in diameter, at the focal plane. Emitted fluorescence is collected and transmitted through the dichroic mirror and an emission filter (Thorlabs, MF530-43) before reaching a 100-mm tube lens (Thorlabs, AC300-100-A) to form an image on a fast scientific CMOS camera (Hamamatsu Photonics, ORCA-Lightning C12120-20P) capable of kHz framerates. The spatial sampling rate of the microscope was calculated as 5.5μm/8× = ∼688nm and experimentally confirmed using a calibration test target. Excitatory power (mW) was measured on a power meter (Thorlabs PM100D, S130C sensor) before each experiment and converted to irradiance (mW/mm^2^) within the excitatory spot area for direct comparisons between datasets.

Time series data was extracted by first motion correcting the videos using NoRMCorre (MATLAB). ROIs were then drawn by hand in Fiji (ImageJ), and an ROI representing the background subtracted from the time series. The background subtracted time series data was then corrected for photobleaching and normalized to 1 by dividing it by a lowpass filtered version of itself using a 0.5- to 2-Hz lowpass second-order butterworth filter. This had the desired effect of capturing small changes due to movement and eliminating them, something a monoexponential fit did not do well. The data was then put through our spike detection algorithm, choosing a minimum of 10 spike templates per cell. To obtain the theta phase, we bandpass filtered the fluorescence traces between 6 and 11 Hz using a first order butterworth filter, giving us the theta frequency band. We then applied the Hilbert transform, giving us the instantaneous amplitude and phase of the bandpass filtered signal. We were able to detect a very consistent relationship between theta phase and spiking across all mice and all indicators, with spike likelihood peaking at 0 radians (**Fig. 4E**). Note that the ASAP3-Kv plot had to be flipped, since it is a downward going indicator and reports all electrical activity in the opposite direction as ASAP4b-Kv and ASAP4e-Kv, meaning it “peaks” in theta pi radians off from the upward indicators unless the response is flipped.

### In vivo 2-P imaging of hippocampus during spatial navigation task

Prior to surgery, imaging cannula implants were prepared using similar methods to those previously published (*31*). Imaging cannulae consisted of a 1.3-mm-long stainless steel cannula (3 mm outer diameter, McMaster) glued to a circular cover glass (Warner Instruments, #0 cover glass 3mm diameter; Norland Optics #81 adhesive). Excess glass overhanging the edge of the cannula was shaved off using a diamond tip file. C57B6/J mice (Jackson Laboratory stock #000664) were first anaesthetized by an intra-peritoneal injection of a ketamine/xylazine mixture. Before the start of the surgery, animals were also subcutaneously administered 0.08 mg Dexamethasone, 0.2 mg Carprofen, and 0.2 mg Mannitol. After one hour, animals were maintained under anesthesia via inhalation of a mixture of oxygen and 0.5–1% isoflurane. Then, a mixture of adeno-associated viruses containing ASAP4b-Kv and jRGECO1b constructs (n = 3 mice; AAV8-syn-ASAP4b-Kv-WPRE, titer=1.17 × 10^12^ vg/mL; AAV1-syn-NES-jRGECO1b-WPRE-SV40, titer = 1.3 × 10^13^, AddGene 100857) was injected into the left hippocampus (500 nL injected at –1.8 mm anterior/posterior [AP], –1.1 mm medial/lateral [ML], 1.4 mm from the dorsal surface [DV]) using a 36-gauge Hamilton syringe (World Precisions Instruments). The needle was left in place for 15 minutes to allow for virus diffusion. The needle was then retracted and the imaging cannula implant was performed.

A 3-mm-diameter craniotomy was performed over the left posterior cortex (centered at -2 mm AP, -1.8 mm ML). The dura was then gently removed and the overlying cortex was aspirated using a blunt aspiration needle under constant irrigation with sterile artificial cerebrospinal fluid (ACSF). Excessive bleeding was controlled using gel foam that had been torn into small pieces and soaked in sterile ACSF. Aspiration ceased when the fibers of the external capsule were clearly visible. Once bleeding had stopped, the imaging cannula was lowered into the craniotomy until the cover glass made light contact with the fibers of the external capsule. In order to make maximal contact with the hippocampus while minimizing distortion of the structure, the cannula was placed at approximately a 10 degree roll angle relative to the animal’s skull. The cannula was then held in place with cyanoacrylate adhesive. A thin layer of adhesive was also applied to the exposed skull. A number 11 scalpel was used to score the surface of the skull prior to the craniotomy so that the adhesive had a rougher surface on which to bind. A headplate with a left-offset 7-mm diameter beveled window was placed over the secured imaging cannula at a matching 10-degree angle, and cemented in place with Met-a-bond dental acrylic that had been dyed black using India ink to prevent VR light from coming into the objective.

At the end of the procedure, animals were administered 1 mL of saline and 0.2 mg of Baytril and placed on a warming blanket to recover. Animals were typically active within 20 minutes and were allowed to recover for several hours before being placed back in their home cage. Mice were monitored for the next several days and given additional Carprofen and Baytril if they showed signs of discomfort or infection. Mice were allowed to recover for at least 10 days before beginning water restriction and VR training.

All virtual reality environments were designed and implemented using the Unity game engine (https://unity.com/). Virtual environments were displayed on three 24-inch LCD monitors that surrounded the mouse and were placed at 90 degree angles relative to each other. A dedicated PC was used to control the virtual environments and behavioral data was synchronized with calcium imaging acquisition using transistor-transistor logic (TTL) pulses sent to the scanning computer on every VR frame. Mice ran on a fixed axis foam cylinder,r and running activity was monitored using a high precision rotary encoder (Yumo). Separate Arduino Unos were used to monitor the rotary encoder and control the reward delivery system.

In order to incentivize mice to run, the animals’ water intake was restricted. Water restriction was not implemented until 10–14 days after the imaging cannula implant procedure. Animals were given 0.8–1 mL of 5% sugar water each day until they reached ∼85% of their baseline weight and given enough water to maintain this weight.

Mice were handled for 3 days during initial water restriction and watered through a syringe by hand to acclimate them to the experimenter. On the fourth day, we began acclimating animals to head fixation (day 4: ∼30 minutes, day 5: ∼1 hour). After mice showed signs of being comfortable on the treadmill (walking forward and pausing to groom), we began to teach them to receive water from a “lickport”. The lickport consisted of a feeding tube (Kent Scientific) connected to a gravity fed water line with an in-line solenoid valve (Cole Palmer). The solenoid valve was controlled using a transistor circuit and an Arduino Uno. A wire was soldered to the feeding tube and capacitance of the feeding tube was sensed using an RC circuit and the Arduino capacitive sensing library. The metal headplate holder was grounded to the same capacitive-sensing circuit to improve signal-to-noise, and the capacitive sensor was calibrated to detect single licks. The water delivery system was calibrated to deliver ∼4 uL of liquid per drop.

After mice were comfortable on the ball, we trained them to progressively run further distances on a VR training track in order to receive sugar water rewards. The training track was 450 cm long with black and white checkered walls. A pair of movable towers indicated the next reward location. At the beginning of training, this set of towers were placed 30 cm from the start of the track. If the mouse licked within 25 cm of the towers, it would receive a liquid reward. If the animal passed by the towers without licking, it would receive an automatic reward. After the reward was dispensed the towers would move forward. If the mouse covered the distance from the start of the track (or the previous reward) to the current reward in under 20 seconds, the inter-reward distance would increase by 10 cm. If it took the animal longer than 30 seconds to cover the distance from the previous reward, the inter-reward distance would decrease by 10 cm. The minimum reward distance was set to 30 cm and the maximal reward distance was 450 cm. Once animals consistently ran 450 cm to get a reward within 20 seconds, the automatic reward was removed and mice had to lick within 25 cm of the reward towers in order to receive the reward. After the animals consistently requested rewards with licking, we began training on tracks used for imaging.

Two visually distinct VR tracks were used for imaging. Each track was 200 cm in length with a 50 cm hidden reward zone. In the first VR track, the 50 cm reward zone was the last 50 cm of the track, and in the second VR track, the 50 cm reward zone began 75 cm down the VR track so that it was in the middle of the track. Mice had to lick within the reward zone in order to receive liquid rewards. At the end of each trial, the animal was teleported to a dark hallway for a randomly determined timeout period of 5-10 sec, chosen with equal probability for each possible timout length. During this timeout period, the laser power was reduced to 0 mW in order to reduce excessive photobleaching. After the timeout period finished, the laser power was increased and the mouse self initiated the beginning of the next trial by running forward. Data from both VR tracks was included in all analyses. To image the calcium and voltage activity of populations of neurons in CA1, we used a resonant-galvo scanning two photon microscope (Neurolabware). All data was collected using a Leica HC IRAPO L 25× objective (1.00 NA, 2.6 mm WD). Neurolabware microscope firmware was modified to allow continuous bidirectional scanning at 989 Hz without digitizer buffer overload (16 × 796 pixels, 0.02 mm × 0.64 mm FOV). 940-nm light (Coherent Discovery laser) was used for excitation of both jRGECO1b and ASAP4b-Kv in all cases. Laser power was controlled using a pockels cell (Conoptics). Laser power was set on each session to allow good SNR with the least amount of power possible. For ASAP4b-Kv imaging, the power ranged from 366 to 645 mW/mm^2^. Light was collected using a Hamamatsu H10770B-40 (green channel) and Hamamatsu H11706-40 MOD (red channel) photomultiplier tubes.

Data was motion corrected using the motion correction pipeline from the Suite2P software package (https://github.com/MouseLand/suite2p). Putative CA1 cell membrane segments were circled by hand using the motion corrected average ASAP4b-Kv image from each session in ImageJ (imagej.nih.gov). Given the dense labeling of cells as well as the dense cell packing of the CA1 pyramidal cell layer, some of the membrane segments likely contain signals mixed from several cells. Based on the numerical aperture of the objective and our approximate axial resolution, we estimate that each ROI contains signal from (∼1–3) cells. Pixel-averaged timeseries were extracted from both the red and green channels for each ROI.

For each ROI, we calculated Δ*F*/*F* independently for the green and red channels. Since laser power was set to 0 mW between each trial, signal baseline was calculated independently on each trial as well. Due to the small FOV and the fact that animals were running at high speeds (∼40 cm/s) some frames were not able to be accurately motion corrected. We attempted to remove the effect of these high-motion frames from our analyses by using Suite2P’s motion estimates to calculate which frames were corrupted by motion. Any frame within 20 frames of one of these high motion frames was replaced with a “NaN” value. Motion estimates from Suite2P rigid motion correction were then used as nuisance regressors for the remaining timepoints, with the following design matrix: *X* = [*x*(*t*), *y*(*t*), *x*^1^ (*t*), *y*^1^(*t*), *x*(*t*)*x*(*t*)]. The residual timeseries from this regression were used for the remaining analyses. For baseline calculation, NaNs were linearly interpolated from the surrounding frames. These motion-imputed timeseries were then low pass filtered to calculate a baseline timeseries (green channel: 0.5 Hz 8^th^ order Butterworth low pass filter, red channel: 0.25 Hz 8^th^ order Butterworth low pass filter). Residual ROI timeseries (NaNs included) were divided by this baseline to get Δ*F*/*F*. For place cell identification and visualization, Δ*F*/*F* was convolved with a 5-frame Gaussian.

Place cells were identified using a previously published spatial information (SI) metric (*43*)

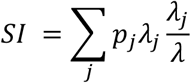

where *λ*_*j*_ is the average activity rate of a cell in position bin *j, λ* is the position-averaged activity rate of the cell, and *p*_*j*_ is the fractional occupancy of bin *j*. The track was divided into 10 cm bins, giving a total of 20 bins.

To determine the significance of the SI value for a given cell, we created a null distribution for each cell independently using a shuffling procedure. On each shuffling iteration, we circularly permuted the cell’s time series relative to the position trace within each trial and recalculated the SI for the shuffled data. Shuffling was performed 100 times for each cell, and only cells that exceeded all 95% of permutations were determined to be significant “place cells”.

To further ensure the reliability of the place cells, we implemented split-halves cross-validation. Taking only the odd-numbered trials, we computed the average firing rate map to identify the position of peak activity. Each cell’s activity was “z-scored” based on the mean and standard deviation across spatial bins on odd-numbered trials. Cells were sorted by this position and then the average activity on even-numbered trials was plotted. This gives a visual impression of the reliability of the place cells. For visualization, single trial activity rate maps were smoothed with a 20-cm (2 spatial bins) Gaussian kernel.

### Gaussian log likelihood probability signal detection technique

The spike detection framework we used builds on the log likelihood ratio based framework created in Wilt. et. al. 2013. To implement the equations developed in Wilt et. al. 2013 on real world, empirical data however, some changes were required. The first was to the underlying assumptions about the noise, where we changed them from theoretical shot-noise limited Poisson assumptions to empirically determined gaussian ones. We found this to be necessary because additional noise is added to the photon-shot noise when the data is collected, and comes from the camera, electric noise in the cables, and many other possible sources. This caused the Poisson assumptions from Wilt et. al. 2013 to often underestimate the actual noise levels in most of our datasets.

Spike templates were chosen for each imaging session, and were chosen to be the largest spiking events from the beginning of the session, with a minimum of 10 chosen each time. Naturally, as photobleaching occurred across trials and spikes become noisier, fewer were detected since the template spikes were taken from the initial imaging period and were thus much larger. While taking only the largest events to use as the template results in very conservative detection, with smaller, noisier events remaining undetected, we wanted to be confident in the events we did detect, and felt that this was a safer approach. Example log likelihood traces are shown in **Fig. 4B** in black. The red line represents the significance threshold, which was chosen such that after breaking the data vector into pieces with lengths equal to the length of the spike template, there was a 1/(20 × number of pieces) chance of getting a false positive in any given piece of data. This was done to correct for multiple comparisons.

Once the event templates were chosen, the equations and procedures described below allow the data to be converted into a probability vector. The probability vector quantifies the probability that the data came from the distribution defined by the templates. N is the length of the templates in samples (time bins), and the data is taken in N samples at a time and compared to it. The background also has a template of length N, but since our data was normalized to 1, we set the background to a vector of ones of length N. See the section below on *Background Noise Estimation* for how we estimated the standard deviation of the background template. Our new gaussian equations for calculating the log likelihood probability that the observed data came from the distribution defined by the template events are as follows:

For time bins where the *k*’th mean template value is less than the *k*’th value of the background, where *k* only counts the time bins where this condition is true. *U* is the total number of time bins where the mean of the templates are less than the mean of the background. *f*_*k*_ is the *k*’th sample of the new data being analyzed.

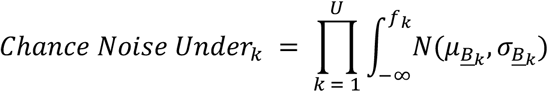

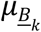 is the mean of the mean of the background template at the *k*’th data point. For our purposes this was the number 1 for all *k* since we normalized the data, but it does not have to be. 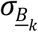 is the standard deviation of the background at the *k*’th point. See the section *Background Noise Estimation* for how this was estimated.

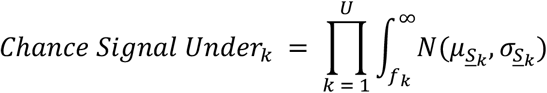

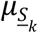 is the mean of the templates the user selected at the *k*’th point. Note that this allows for the shape of the event to be taken into account when calculating probabilities. 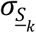 is the standard deviation of the *k*’th time bin for the templates chosen. This means that each time bin of the event template has its own mean and standard deviation, creating N gaussian distributions, which are needed to take the gaussian integrals at each time bin. N is the length of the templates in time bins.

Below, for time bins when the *j*’th mean template value is greater than the *j’*th value of the background, where *j* only counts the time bins where this condition is true. *O* is the total number of time bins where the mean of the templates are greater than the mean of the background. *f*_*j*_ is the *j*’th sample of the new data being analyzed.

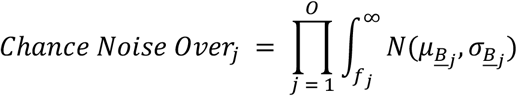

This is the same as above, only for time bins where 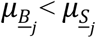, so the limits of integration change.

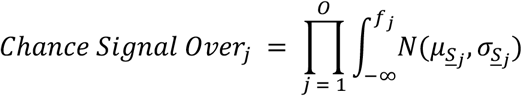

This is the same as above, only for time bins where 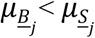, so the limits of integration change.

These probabilities are multiplied together, the log of the ratio of them obtained, and ultimately a single number for the *i’*th sample of the data vector is returned, where *i* runs from 1 to the length of the data vector.

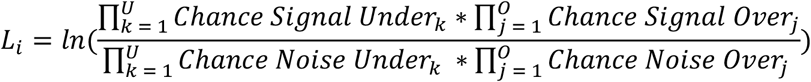

Note that since N is the length of the template in samples, N = *U* + *O*.

This is repeated, each time shifting over by a single time bin in the data, until the entire data vector has been analyzed and converted into a probability vector. The very end of the data vector is chopped as we did not find an objective way to pad the end of data vector such that it would not impact the probabilities calculated for when the template distribution reaches the end. Fortunately, our templates were typically very short in length, so this resulted in a very negligible loss of data at the very end of each data set (a few tens of milliseconds). While taking the log of the ratio here is not strictly necessary for the purposes of signal detection, it helped to keep the ratio from becoming too positive or too negative to visualize effectively when making plots.

We next find all of the events that cross the probability threshold we set. Every segment above threshold is considered a single template matching event, for as long as it stays above threshold in the log likelihood probability vector. Since it does this, it is important to set the filter parameters such that you pull out the events you care about. For example, if the user wanted to detect single spikes riding on top of depolarizing calcium waves, as well as the depolarizing waves themselves, you would first set a very low frequency lowpass filter as the baseline calculation, and choose the entire duration of the burst as a template. Once you have the detected bursts pulled out, you would then set the passband of the lowpass filtered calculation of the baseline higher than what was used previously to pull out the entire burst, in order to flatten out the calcium portion of the event, while maintaining the faster spikes for detection. This enables the user to sequentially detect bursts, followed by spikes within bursts.

For onset timing, the time point where the log likelihood ratio first crosses the threshold is considered the start of the event. For the event detection performed on the 1-P in vivo imaging dataset of SOM+ interneurons, we first highpass filtered both the spike templates and raw data with a 20-Hz second-order Butterworth filter.

### Background noise estimation

The standard deviation of the background template in this case is assumed to be constant across the background template. It is recalculated for each new batch of data that is fed into the program, and involves 3 successive attempts, if the previous attempt fails.

First attempt: The program fits a gaussian mixed model (GMM) to the entire data vector, with the assumption that there are two gaussians present. One is the signal, one is the noise. This tends to work well for traces where lots of activity is present, giving the histogram of the data vector a tail to one side; otherwise it tends to only find a single gaussian. The standard deviation of the noise is set to be the std. of the gaussian on the left if the indicator is upward going, or the gaussian on the right if it is downward going, which the program detects from the peak of the average templates. This prevents overestimating the noise, which would result in more false negatives than desired (but fewer false positives as well).

Second attempt: If the standard deviation of the GMM fit is greater than the standard deviation of the whole data vector, then something bad happened with the fit because that doesn’t make any sense. The program then reverts to what we decided to call the “asymmetric distribution technique”.

Asymmetric distribution technique: If the data contains signals, the distribution of the data will be skewed to one side. To the right if the indicator is upward going, and to the left if it is downward going. The side of the distribution away from the direction the indicator moves in must be pure noise, with no contamination from the signal we are looking for. That half of the distribution is then flipped about the mean to replace the half that is a mix of noise and signals, giving a good estimation of how the distribution of the data would look if there were no signals present in it. The standard deviation of this “flipped” distribution is then taken to be the standard deviation of the noise.

Third attempt: Occasionally, attempt 2 also fails, possibly due to movement or lots of hyperpolarizing activity, and the standard deviation of the flipped distribution is greater than the standard deviation of the data itself, which again does not make sense. This happens very rarely, but in this case the standard deviation of the background is simply assigned the value of the standard deviation of the entire data vector. This will have the effect of giving too many false negatives, but it has the advantage of giving fewer false positives as well, so represents a conservative estimate of spike probability.

Note that how the background noise is chosen will effect the numbers in the log likelihood vector. Smaller background noise values will result in larger log likelihood values, and vice versa. This is because that as the noise distribution gets narrower via a smaller standard deviation, the probability that the same data point came from the noise distribution shrinks.

The code was written such that if there are more than 9 templates, the standard deviation of the templates themselves are taken, as described previously, to be the standard deviation of the templates at each time bin in the template vector. If there are less than 10 templates, the std of the templates is assumed to be the same as the noise, and only the mean changes (the mean template trace is still taken, each time bin just has the same std which is the same as the noise).

## ACKNOWLEDGEMENTS

We thank Dario Ringach (Neurolabware) for help with firmware changes to the 2-P microscope used for in vivo hippocampal imaging in the Giocomo lab. We thank Helen Yang and Marjorie Xie for the stimulus generation and data analysis code used in the Clandinin lab. We thank Celina Yang and Max Melin for mouse surgical preparations for 1-photon hippocampal imaging in the Golshani Lab.

## Author Contributions

Experimental design and study conception was done by M.Z.L., S.W.E., A.G., J.T, B.M., S.C. VK. and M.P. All cell culture screening was done by D.S., M.C, S.W.E. and L.P. All cell culture patch clamping was done by D.S.,M.C., D.J. and G.Z. All Acute striatal slice brightness work was done by S.W.E. Hippocampal acute slice work was done by C.D.M., A.R., FJ.H. and S.W.E. All in vivo 2-P work was done by M.P, with feedback from L.M.G. A.N. And A.L. helped with testing ASAP4b and ASAP4e in the hippocampus in vivo. 1-P in vivo work was done by B.M. and J.T., with advice from P.G. and data analysis by S.W.E. All photobleaching and 2-P spectra experiments were done by S.W.E. and S.L. The spike detection work was done by S.W.E. The in vivo ASAP4e and jRGECO1a recordings in vivo were done by J.L.F. in the lab of N.J. All modeling work was done by S.C.VK., and CM.S., with input from R.O.D. Figure prep was done by S.W.E., M.Z.L., G.Z., D.S., M.C., C.D.M., A.R. and M.P. Manuscript writing was done by S.W.E. and M.Z.L. Y.H. assisted with the data analysis of the in vivo cortical pyramidal cell data. All authors reviewed the manuscript.

